# *Toxoplasma* surface coat protein TgSRS57 (TgSAG3) regulates complement deposition, Factor H recruitment, and promotes parasite persistence

**DOI:** 10.1101/2022.10.17.512461

**Authors:** Viviana Pszenny, Patricia M. Sikorski, Raghavendran Ramaswamy, Michelle L. Parker, Gloria Adedoyin, Martin J. Boulanger, Michael E. Grigg

## Abstract

The surface coat of *Toxoplasma gondii* possesses a lectin-like activity that activates host complement. Serum resistance relies on specific recruitment of host regulators Factor H (FH) and C4BP to the parasite surface to inactivate bound C3 and evade the complement system. TgSRS57 (TgSAG3) was previously reported to possess a lectin-like activity specific for sulfated proteoglycans (SPGs) that facilitated host cell attachment and parasite invasion. To investigate whether TgSRS57 actively recruits Factor H to the parasite surface to regulate the complement cascade, we generated Type I RH and Type II CZ1 knockout strains. Δ*srs57* parasites showed a reduction in both C3 deposition and FH recruitment indicating that TgSRS57 is a C3 acceptor and regulates complement activation. However, Δ*srs57* parasites showed no attachment defect, possessed normal extracellular survival and intracellular replication kinetics, and were more virulent than WT strains in low dose murine infections, contrary to previous findings. To better understand the genetic and cellular data, we determined the 1.59Å resolution TgSRS57 crystal structure. Notably, TgSRS57 did not possess a dimer dependent basic surface groove as originally modeled, nor did it bind heparin. However, approximately 10 distinct parasite proteins were identified that interact specifically with SPGs. Infection studies showed that Type II Δ*srs57* parasites achieved higher parasite burdens and elevated inflammatory responses compared to WT parasites. Infection of C3 deficient mice established that TgSRS57-dependent protection was C3-dependent and that *T. gondii* actively parasitizes the complement system. Our results suggest that the interaction between TgSRS57 and C3 plays a critical role in disease tolerance by controlling both parasite proliferation and persistence *in vivo*.

**AUTHOR SUMMARY:** The *Toxoplasma gondii* surface coat is dominated by a superfamily of developmentally expressed, antigenically distinct adhesins collectively called the SRS proteins, which are structurally analogous to the *Plasmodium* 6-CYS surface proteins. These proteins are thought to possess lectin-like activities, facilitate parasite attachment and entry into host cells, and regulate innate effectors of the host immune response. Disease tolerance in hosts is critically dependent on the induction of sufficient immunity to limit parasite replication and promote host survival without inducing bystander immune damage. In this study we employed a combination of genetic, cellular and structural biology approaches to establish that the surface antigen TgSRS57 regulates complement C3 deposition on parasite surfaces by specifically recruiting host Factor H to actively resist serum killing and limit the formation of the C5b-9 membrane attack complex. Further, that the presence of C3 was host protective, as C3-deficient mice failed to control parasite replication. The ability of the parasites to inactivate C3 was also necessary to resist serum killing and promote chronic infection. Because complement exists as a potent inflammatory mediator, the active regulation of complement by *Toxoplasma* ensures that the parasite is not lysed by C3 but also limits the induction of lethal immunopathology, thereby promoting an immune balance, disease tolerance, and the long-term persistence of transmissible parasites.

## INTRODUCTION

The protozoan parasite *Toxoplasma gondii*, the etiological agent of toxoplasmosis, causes significant morbidity and mortality in humans and animals worldwide [1]. In immunocompetent adults, acute *T. gondii* infections can produce influenza-like symptoms before resolving into a life-long infection after the parasite encysts in the muscles and brain. *Toxoplasma* is the leading cause of infectious uveitis globally, with rates of ocular toxoplasmosis ranging from 1-20% among seropositive individuals [2, 3]. *T. gondii* is an AIDS defining opportunistic pathogen, and can produce serious disease in cancer and organ transplant patients [4] where prognosis is poor because no sterilizing prophylactic drugs exist.

*T. gondii* is an obligate intracellular pathogen that must attach to and enter host cells to be infection competent. It relies on a series of molecular interactions that initially “sense” cells [5, 6]. This involves a series of reversible attachments and perturbations of host cell membranes, referred to as the “kiss and spit” model [7], which allows the parasite to identify permissive host cells prior to invasion. The remarkable ability of *T. gondii* to infect essentially any cell type in a broad range of hosts suggests that attachment relies on the ability of the parasite to recognize a series of conserved, ubiquitously displayed surface molecules, or constituents of the extracellular matrix that associate specifically with host cell surfaces. In fact, numerous microbial pathogens have evolved surface coats that recognise and interact with a variety of sulfated glycosaminoglycans to establish infection [8]. Previous work showed that *T. gondii* possesses carbohydrate binding activities that promote the agglutination of erythrocytes, an effect reversed by the addition of sulfated glycoconjugates such as heparin, fucoidan and dextran sulphate. Further, cells deficient in surface proteoglycans, or their sulfation, are more resistant to *T. gondii* infection [6, 9-11]. Collectively, these studies suggest that lectin-like surface protein(s) on *T. gondii* facilitate the recognition of invasion permissive host cells. In addition to the micronemal proteins [6, 12, 13], likely candidates that possess carbohydrate-binding activity are a superfamily of developmentally expressed, antigenically distinct surface coat proteins known as the SRS (SAG-Related Sequence) adhesins [14]. *Toxoplasma* encodes more than 180 SRS adhesins, each possessing a signal peptide and glycosylphosphatidylinositol (GPI) anchor [15, 16]. While the SRS proteins are best studied in *T. gondii* [17, 18], other tissue dwelling coccidia express these genes [19]. Recent work in *P. falciparum* showed that the 14 member 6-Cys family of *Plasmodium* surface proteins, which facilitate invasion of hepatocytes (P52/P36), promote male/female gamete recognition and fertilization (Pfs230, 47, 48/45), CD36 binding (Sequestrin), and the recruitment of host complement regulators (Pf92) are structurally related to the SRS proteins [20-24].

Two *T. gondii* surface antigens, TgSRS29B (formerly SAG1) and TgSRS57 (formerly SAG3), have previously been shown to possess lectin-like activity by their ability to bind sulfated proteoglycans (SPGs) [25, 26]. Whereas the TgSRS29B-deficient mutant shows no appreciable deficit in host cell infection [27], the TgSRS57 deficient parasites were significantly impaired in host cell attachment and murine infection competency [28]. Follow-up binding studies demonstrated a specific interaction between TgSRS57 and heparin [26]. The crystal structure of TgSRS29B rationalized a structural basis for these observations, by revealing a dimer-dependant basic groove that was ideally suited to coordinate a negatively charged host cellular ligand, such as heparan sulfate [29]. Intriguingly, the model developed for TgSRS57 based on the TgSRS29B structure predicted a substantially more basic groove, consistent with its predicted role as a major surface adhesin capable of recognizing negatively charged SPGs [29].

Previous work demonstrated that the surface of *T. gondii* is covalently modified by complement C3b which is rapidly inactivated after exposure to non-immune human serum [30, 31]. The recruitment of soluble host regulators Factor H (FH) and C4b-Binding Protein (C4BP) to the parasite surface coat are significantly associated with *T. gondii* serum resistance *in vitro* [31, 32]. FH is an important negative regulator of the alternative pathway that possesses 20 homologous domains, known as short consensus repeats (SCRs) that bind C3b, sialic acid and SPGs [33, 34]. The *T. gondii* protein(s) that recruit FH remain enigmatic, but recent evidence from *Plasmodium* identified the specific recruitment of FH by a 6-Cys protein Pf92 [22] suggesting that SRS proteins may be involved.

This study was performed to determine if TgSRS57, which is predicted to bind SPGs, is covalently modified by C3 and specifically recruits FH to evade the complement system. To answer these questions, we employed a combination of genetic, cellular and structural biology approaches to show that TgSRS57 regulates C3 complement deposition and FH recruitment to inactivate complement. We found no evidence that it possesses a lectin-like activity. Further, TgSRS57 is an important virulence factor that has no obvious defect in the attachment and invasion of host cells, but rather, it recruits FH to inactivate C3b and induces a protective immune response that regulates both the proliferation and persistence of parasites *in vivo*. Our work also resolved several parasite immunogenic proteins that interact specifically with SPGs and demonstrated that SPGs are required for optimal parasite invasion and growth.

## RESULTS

### Targeted deletion and complementation of TgSRS57 in Type I RH and Type II CZ1

Previous studies suggested that recognition of cellular SPGs by the *Toxoplasma* surface coat protein TgSRS57 promotes parasite attachment, host cell infectivity, and murine infection competency [26, 28]. Furthermore, the prediction that TgSRS57 possesses a positively charged groove (model based on the 1.7 Å resolution TgSRS29B structure) suggested that it was able to accommodate heparin binding without any spatial or steric conflict [29]. These facts prompted us to determine the precise identity of the biologically relevant SPG and the molecular basis underlying infection competency thought to be mediated by a TgSRS57–SPG interaction. In order to address this question, a *Δsrs57* deficient mutant was generated in the RH Type I and CZ1 Type II genetic background. The genomic locus of *TgSRS57* was first disrupted by homologous recombination using an HXGPRT mini gene as the positive selectable marker in the RHΔ*hxgprt* strain **(Fig. 1A)**. Ten clones lacking TgSRS57 expression were identified by screening three independent transfections using an anti-TgSRS57 monoclonal antibody (1F12, a kind gift from Dr. Stan Tomavo) by flow cytometry. Southern blot analysis confirmed the presence of only one insertion in 3 independently derived clones that were selected for further studies (data not shown). Additionally, parasite lysates from these clones were treated with GPI-PLC to expose the cross-reacting determinant (CRD) present in surface-associated SRS proteins that possess a GPI-anchor. GPI-PLC treated lysates were subjected to SDS-PAGE, transferred to nitrocellulose and probed with anti-CRD antibodies. Consistent with the result of the flow cytometry analysis, the band corresponding to TgSRS57 was absent in the three Δ*srs57* clones (**Fig. 1B**-lanes 3, 4 and 5). Moreover, there did not appear to be any compensatory change in the surface expression of other GPI-linked SRS surface coat proteins (**Fig. 1B**). In order to restore the expression of TgSRS57 at a heterologous site within the *T. gondii* genome, and to generate transgenic parasites expressing the green fluorescence protein (GFP) and firefly luciferase protein (LUC) for *in vivo* imaging experiments, Δ*srs57* parasites (as well as the previously published French RH (fRH) Δ*srs57* mutant CL55, kindly provided by Dr. S. Tomavo) were co-transfected with a 6.7kb PCR amplified genomic locus of Tg*SRS57*, encompassing the 5’ and 3’ UTRs and the promoter region as well as a pGFP-LUC plasmid, containing cDNA sequences of GFP and LUC. The strategy for complementation is depicted in **Supplemental Fig. 1**. After transfection, parasites positive for GFP were sorted by flow cytometry and GFP positive clones were assessed for surface expression of TgSRS57. No difference was detected in the expression of TgSRS57 between wild type (WT) parent and the complemented clones by flow cytometry (**Fig. 1C**-(left)). The expression of TgSRS34A, a highly immunogenic surface antigen, was unchanged in the knockout and complemented parasites (**Fig. 1C** (right)).

**Figure 1.**
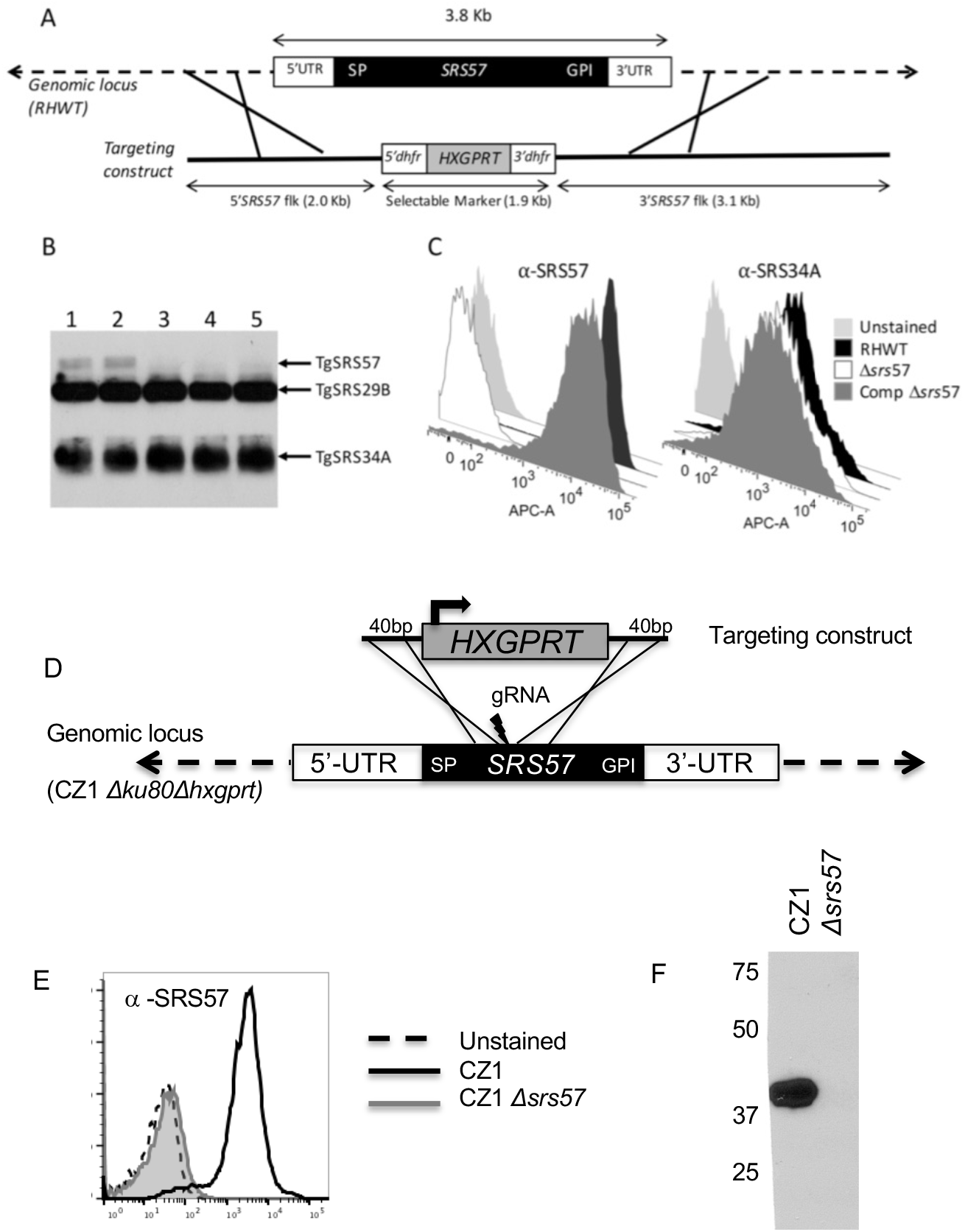
Targeted deletion of TgSRS57 in Type I RH and Type II CZ1. **(A)** (Top) Diagram of a genomic fragment spanning the TgSRS57 genomic locus. The TgSRS57 coding sequence is indicated by a black box. SP indicates the signal peptide and GPI the glycosylphosphatedylinositol anchor. 5’ and 3’ UTRs are indicated by white boxes. (Bottom) 5’dhfr-HXGPRT-3’dhfr construct used to disrupt the TgSRS57 gene by homologous recombination. **(B)** Anti-CRD Western blot of RHWT (lane 1), RH TgSRS57 mistargeted (lane 2) and 3 independent clones of RHΔ*srs57* strain (lanes 3, 4 and 5), showing the absence of TgSRS57. Tachyzoites pellets of each strain, were solubilized in 1% NP-40, treated with GPI-PLC, separated by SDS-PAGE, transferred to nitrocellulose and probed with anti-CRD polyclonal antibody. **(C)** Flow cytometry analysis of RHWT (black), RHΔ*srs57* (white) and complemented (dark grey) strains labelled with anti-TgSRS57 (1F12 monoclonal antibody) on the left and anti-TgSRS34A (5A6 monoclonal antibody) on the right. Expression of TgSRS57 was completely abolished in the knockout strain, whereas the expression of TgSRS34A was not affected. **(D)** Schematic representation of knockout construct for TgSRS57. The endogenous locus was disrupted using the hypoxanthine-xanthine-guanine phosphoribosyl transferase (HXGPRT) selectable marker and CRISPR methodology. **(E)** Flow cytometry analysis of CZ1 wild type (black) and Δ*srs57* (gray) labeled with anti-TgSRS57 (1F12 monoclonal antibody). **(F)** Western blot analysis of 1×10^6^ CZ1 wild type strain and Δ*srs57* parasites. Nitrocellulose membranes were probed with anti-TgSRS57 (1F12 monoclonal antibody).

To generate a Δ*srs57* clone in the CZ1 Type II strain, we took a CRISPR/Cas9 approach to disrupt the *TgSRS57* locus by inserting an *HXGPRT* mini gene after the *TgSRS57* signal peptide in a recently generated CZ1Δ*ku80*Δ*hxgprt* strain. A single guide RNA was designed to introduce a break within the 5’ end of the *TgSRS57* locus. The *HXGPRT* gene was amplified using primers that possessed 40bp of homology with the *TgSRS57* gene either side of the guide RNA cut site (**Fig. 1D**). After transfection and drug selection, screening for independent clones deficient in TgSRS57 was done by PCR. Clones positive for *HXGPRT* at the *TgSRS57* locus were then tested for lack of TgSRS57 protein expression using the anti-TgSRS57 monoclonal antibody (**Fig. 1E**) and by Western blot (**Fig. 1F**).

### Parasites lacking TgSRS57 display normal extracellular survival kinetics

To determine if the lack of TgSRS57 impacted parasite extracellular survival, a time course experiment was performed. Parental, Δ*srs57* and complemented strains were subjected to extracellular conditions during 0, 0.5, 1, 2, 4, 8, 24 and 48 hours prior to infecting a confluent monolayer of human foreskin fibroblasts (HFF) cells with a dose of 10^3^ tachyzoites/strain/time point. Analysis of the linear regression slopes showed no significant differences between the three strains, indicating that the deletion of TgSRS57 had no impact on extracellular persistence **(Fig. 2A**). Comparable results were obtained with the French RHΔ*srs57* mutant CL55 and its complemented control (**Supplemental Fig. 2**). Similarly, a paired comparison between RH versus French RH strains (either wild type, knockout or complemented) showed no significant differences regardless of the strain investigated (**Supplemental Fig. 2**). Time points of 24 and 48 hours were not presented because no plaque formation was observed, indicating that the parasites had a finite time in which they survived outside the host cell.

**Figure 2.**
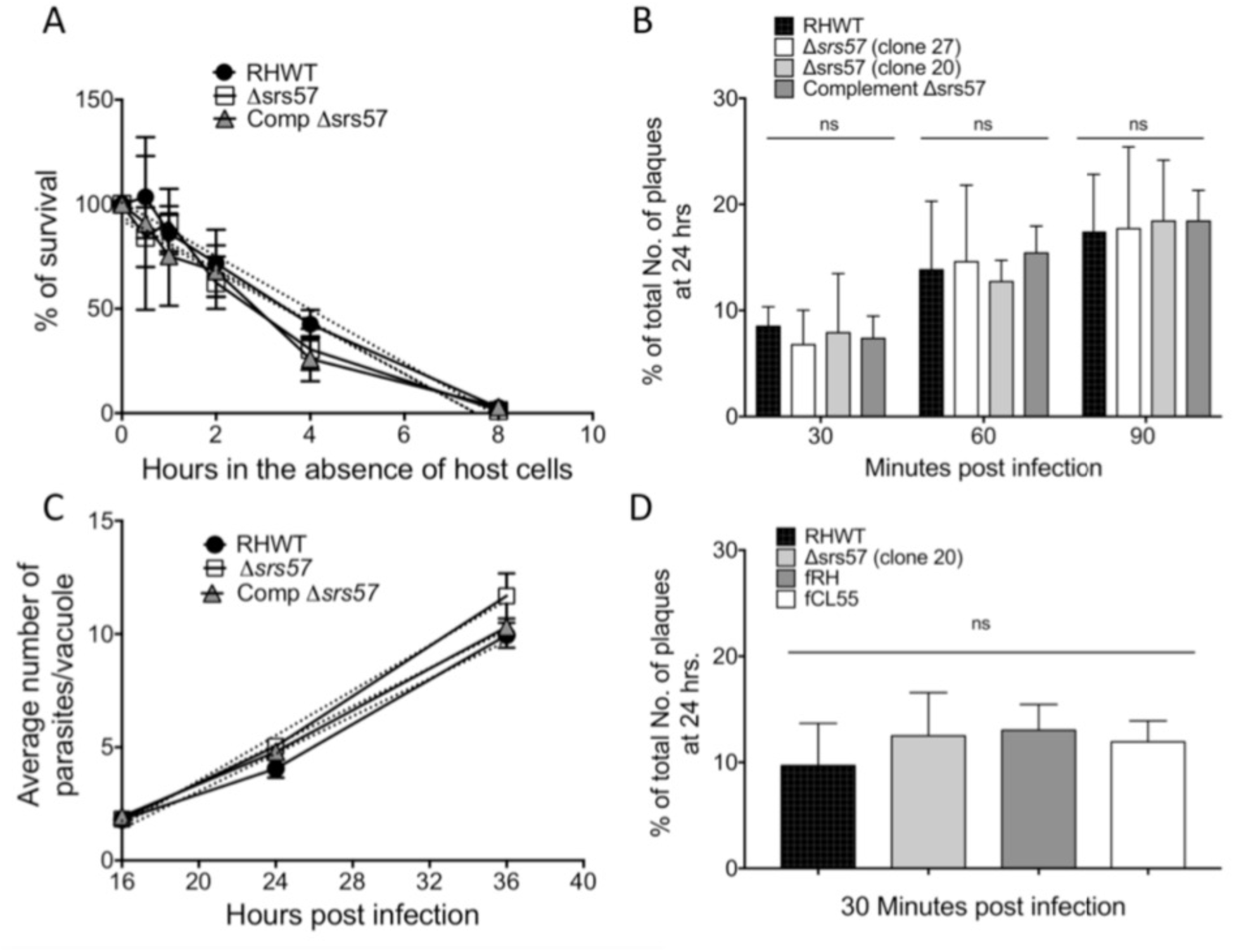
In vitro characterization of extracellular survival, attachment, invasion, plating efficiency and intracellular growth of Δsrs57 parasites. **(A)** Quantification of the extracellular survival. 10^3^ tachyzoites of each strain (RH WT, Δ*srs57* and complement) were maintained for 0 min, 30 min, 1, 2, 4 and 8hrs in extracellular conditions, before applying to a fresh monolayer of HFF cells. After 5 days in normal culture conditions, the number of plaques formed at each time point were counted. No significant differences were observed in the extracellular survival and plating efficiency between the strains. Values represent the mean of triplicates ± SD. **(B)** Time course plating efficiency. 10^3^ tachyzoites of each strain (RH WT and Δ*srs57* [clones 20 and 27], and complemented Δ*srs57*) were inoculated onto monolayers of HFF cells, grown in 24 well plates, and allowed to invade for 30, 60, 90 min, and 24 hrs. After extensive washes, the parasites were incubated undisturbed in normal culture conditions and the total number of plaques were recorded after 6 days. The results were expressed as a percentage of total number of plaques counted at 24 hrs. Data represent the mean of 3 independent experiments ± SD. The statistical significance was analyzed by the One-way ANOVA (non-parametric) test, using Prism 7 software. The results show no significant differences between strains, indicating that the lack of TgSRS57 does not affect the ability to invade. **(C)** Rate of replication. Confluent monolayers of HFF cells, were infected with 10^5^ tachyzoites/well of each strain and incubated for 16, 24, and 36 hrs. The average number of parasites/vacuole was scored by counting 100 randomly selected vacuoles. The results are presented as the mean of two independent experiments ± SD. No significant differences in *in vitro* intracellular proliferation were observed between RHWT, Δ*srs57* and the complemented strains. **(D)** Comparison of plating efficiency between RH and fRH strains. No significant differences in plating efficiency were observed between strains. 10^3^ tachyzoites of each strain (RH WT, Δ*srs57* [clone 20], fRH and fRH*Δsrs57* [clone CL55]) were inoculated onto monolayers of HFF cells grown in 12 well plates, allowed to invade for 60 min, washed extensively, and incubated undisturbed in normal culture conditions. The total number of plaques was recorded after 6 days. The results were expressed as a percentage of total number of plaques counted at 24 hrs. Data represent the mean of 3 independent experiments ± SD. The statistical significance was analyzed by One-way ANOVA (non-parametric) test using Prism 7 software.

### *TgSRS57* deficient parasites show no attachment defect in a time course plaque assay

Previous work suggested that Δ*srs57* mutants were defective in their ability to enter HFF cells [28]. Plating efficiency was thus determined using a time course plaque assay by inoculating HFF cells with 10^3^ tachyzoites of each strain (RH, Δ*srs57*, and complemented control). After 30, 60, and 90 minutes the monolayers were extensively washed to remove uninvaded parasites. To determine the total number of invasion events possible in a 24 hour period (to serve as denominator for each individual strain tested) 250 tachyzoites were allowed to enter HFF cells. After 6 days, the number of plaques were counted and the percentage of plaques that formed compared to the 24-hour time point was calculated. No significant difference was observed between all strains tested (**Fig. 2B)**. Our findings were not in accordance with previously published work that used the French RH strain (referred to as fRH, a Type I strain that is genotypically different from the RH strain isolated in the USA by Sabin in 1939). To determine if this was a peculiarity of the mutant clone generated using the fRH strain, we assayed the Δ*srs57* CL55 mutant and its fRH parent. No difference was detected between fRH, the CL55 Δ*srs57* clone and RH and Δ*srs57* clone 20 generated above for 30 (**Fig. 2D**), 60 or 90 minutes (data not shown) in the time course plaque assay performed as in Figure 2B. When the complemented CL55 clone, which expressed wild type levels of TgSRS57, was tested against RH and the Δ*srs57* clone 20, no defect in extracellular survival was detected *in vitro*. (**Supplemental Fig. 2**).

### Intracellular replication is not affected by lack of TgSRS57

To assess if there was any defect in the rate of intracellular growth in HFF monolayers, the number of parasites per vacuole were counted after 16, 24 and 36 hours post-infection. As determined for extracellular survival, the rate of growth did not differ significantly in the Δ*srs57* parasites compared with the parental wild type and complemented strains **(Fig. 2C)**.

### TgSRS57 regulates C3b deposition

Previous work demonstrated that *T. gondii* encodes surface coat proteins that are covalently opsonized by the complement effector protein C3b, which gets rapidly inactivated (within minutes) to limit MAC complex formation and confer resistance to lysis in non-immune human serum [30, 31]. A flow cytometry-based C3b deposition time course assay was performed on parasites incubated in the presence of 10% non-immune human serum (NHS). Consistent with our previous study, Type II strains showed significantly higher levels of C3 deposition compared to Type I strains. However, Δ*srs57* parasites, in comparison to their respective Type I or Type II WT stain, showed a significant (∼50%) reduction in C3b deposition. These results suggest that TgSRS57 is either a parasite surface coat protein that gets covalently modified by C3b or limits its deposition on the parasite surface (**Fig. 3A, 3B)**. Western blot analysis of C3 activation and inactivation products identified iC3b degradation products at 68 and 43 kDa within 10 minutes in both WT and *Δsrs57* parasites, consistent with host mediated inactivation of C3b by Factor H and serine protease Factor I **(Fig. 3E**).

**Figure 3.**
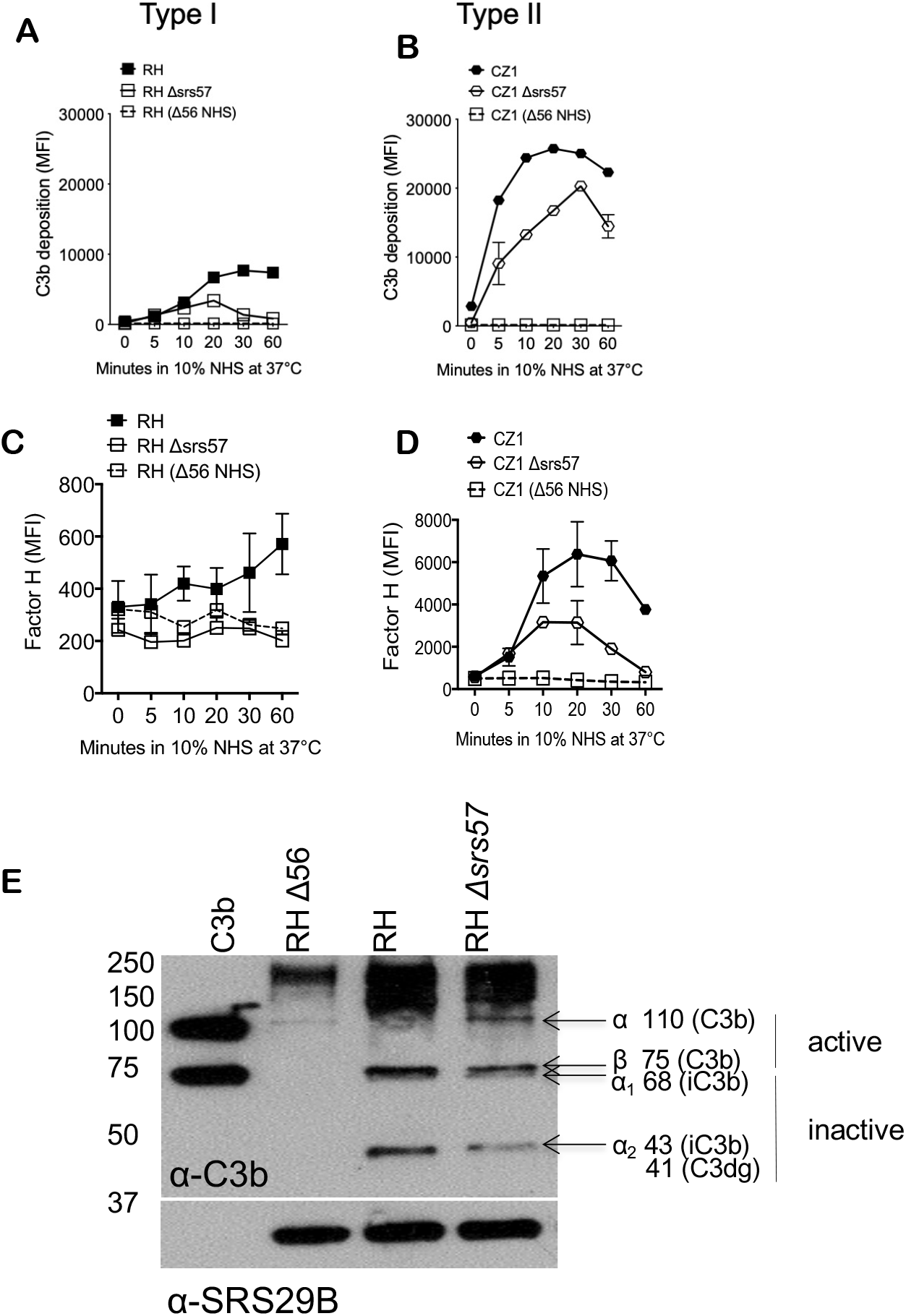
TgSRS57 impacts complement deposition. C3b deposition time course of **(A)** RH wild type (black square) and RH *Δsrs57* (open square) and **(B)** CZ1 (black hexagon) and CZ1 *Δsrs57* (open hexagon) incubated in 10% non-immune human serum (NHS) over 60 minutes at 37°C. Serum was diluted in complement activation buffer, Hanks Buffered Saline Solution supplemented with 0.15mM CaCl_2_ and 1mM MgCl_2_ (HBSS_++_). Wild type parasites in 10% heat inactivated serum (Δ56) were used as a negative control (open square, dotted line). C3 deposition was measured by staining parasites with a monoclonal mouse α-human C3b/iC3b antibody (1:500, Cedarlane). Flow cytometry analysis of Factor H recruitment on **(C)** Type I RH (black square) and RH *Δsrs57* (open square) or **(D)** Type II CZ1 (black hexagon) and CZ1 *Δsrs57* parasites incubated in 10% NHS over 60 minutes at 37°C. Wild type parasites in 10% heat inactivated serum (Δ56) were used as a negative control (open square, dotted line). Factor H was measured by staining parasites with goat anti-human Factor H (1:2,000, Complement Technologies). **(E)** Western blot analysis of C3b deposition on RH and RH *Δsrs57* parasites incubated in 10% NHS for 20 minutes at 37°C. Membranes were probed with goat anti-human C3b at (1:20,000, Complement Technologies). Error bars represent mean ± SEM. Time course experiments and western blots are representative of at least two experiments.

### TgSRS57 is required to recruit FH to parasite cell surface

We recently showed that opsonized parasites recruited both Factor H (FH) and C4b-binding protein (C4BP) to inactivate C3b to promote serum resistance [31]. We determined FH was responsible for the majority of this resistance to serum killing. FH recruitment to the surface of host cells is stabilized by binding both C3b and SPGs in close proximity [34] therefore we postulated that TgSRS57, which was shown previously to bind SPGs, recruits FH in an SPG- and C3b-dependent manner. Factor H recruitment was measured by flow cytometry. RH*Δsrs57* parasites largely phenocopied the heat inactivated serum control (Δ56 NHS) indicating that, in the RH background, tachyzoites did not accumulate appreciable levels of FH on their surface (**Fig. 3C**). The Type II CZ1 strain showed greater overall levels of FH on the WT parasite surface compared to RH, and the CZ1*Δsrs57* parasites exhibited a partial reduction (∼50%) in the recruitment of FH (**Fig. 3D**). These results suggest that TgSRS57 plays a role in the binding of host co-factors that regulate C3b deposition, and that its function may be more relevant for Type I strains, than for Type II strains. Further, that Type II strains possess at least one other factor capable of promoting FH recruitment to the parasite surface.

### TgSRS57 lacks the structural hallmarks of an SPG binding protein

To determine precisely how TgSRS57 coordinates a predicted SPG interaction, its structure was determined using X-ray crystallography. TgSRS57 was produced recombinantly in insect cells and purified to homogeneity by nickel affinity and size exclusion chromatography (SEC) (**Fig. 4A**). The protein eluted as a monomer from SEC and migrated as a doublet on SDS-PAGE possibly due to O-linked glycosylation. TgSRS57 crystallized in space group P2_1_2_1_2_1_ with one molecule in the asymmetric unit and the final structure refined to a resolution of 1.59 Å (**Table S1**). TgSRS57 was modeled from Tyr57 to Pro330, with an additional five residues at the C-terminus derived from the expression construct, and two disordered residues in an apical loop (Leu102 and Gly103). Overall, TgSRS57 adopted a dumb-bell shaped architecture with 2 disulfide bond-stabilized β sandwich domains connected by a proline rich linker **(Fig. 4B, inset)**. The D1 domain extended from Tyr57 to Glu198 and formed a five on four β sandwich, while D2 extended from Met204 to Val335 and formed a four on three β sandwich.

**Figure 4.**
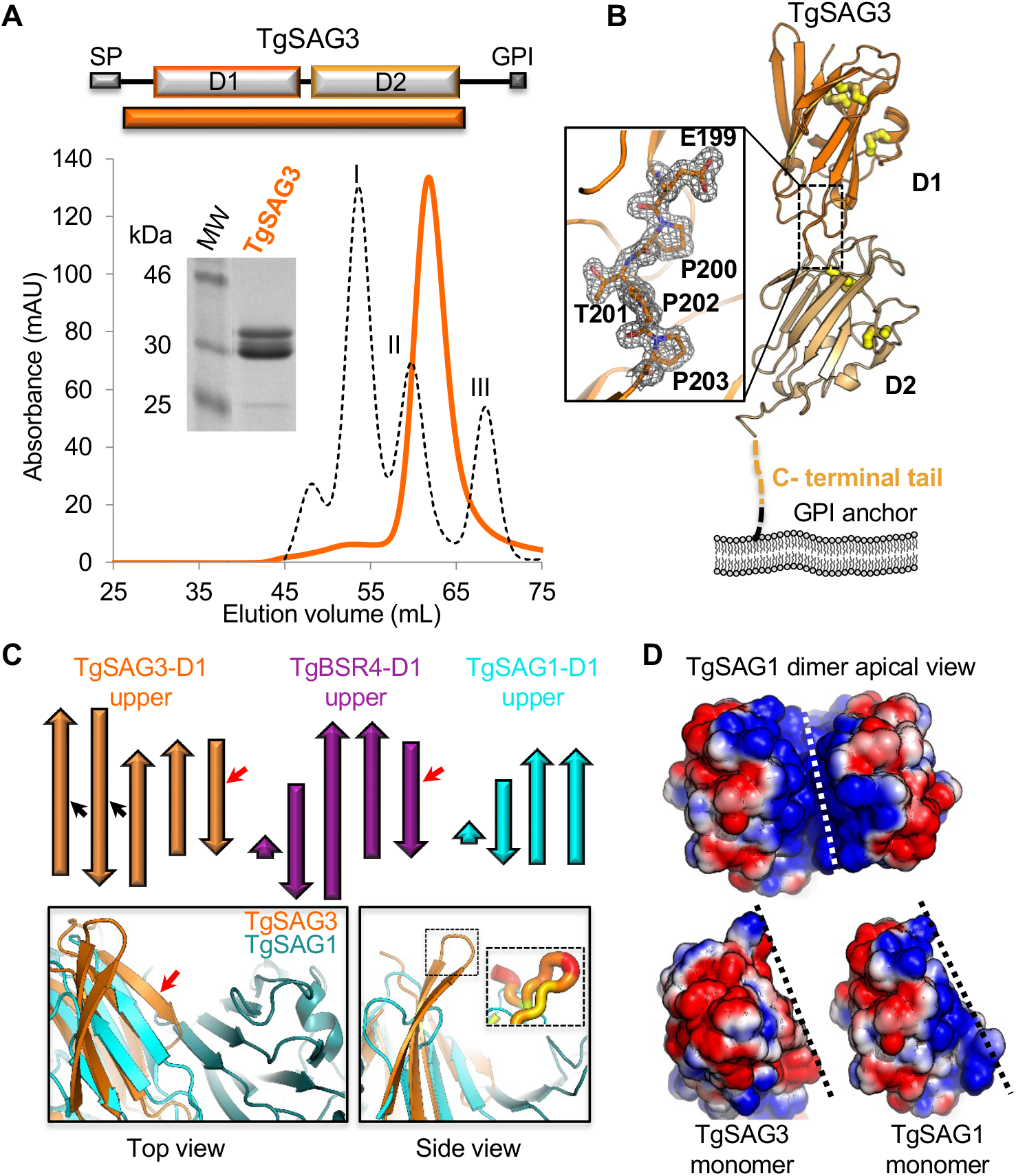
Structural divergence of the SRS fold and biological implications of the TgSRS57 crystal structure. **(A)** Top: Schematic representation of TgSRS57 with predicted structural domains, GPI regions and signal peptides (SP). Orange rectangle represents the crystallization construct. Below-SEC profile of TgSRS57 (orange line). Inset, Coomassie-stained SDS-PAGE gel of TgSRS57 (expected molecular weight: 32 kDa). The protein standards used were conalbumin (75 kDa; peak I), ovalbumin (43 kDa; peak II), and carbonic anhydrase (29 kDa; peak III. **(B)** Left: Tertiary structure showing the head to tail structure of TgSRS57 in the predicted orientation with respect to the parasite cell surface with the N-terminal domain (D1) in bright orange and the C-terminal domain in light orange. The disulphides are indicated as yellow sticks. The C-terminal tail is indicated in light orange. Inset: 2*F*_*o*_ − *F*_*c*_ σA weighted electron density maps at 1.2σ showing the inter-domain linker region. **(C)** Top: Topology diagrams of the β strands that form upper leaves of D1 in TgSRS57 (orange), TgSRS16C (purple), and TgSRS29B (cyan). TgSRS57 is most similar to TgSRS16C and notably more substantial than TgSRS29B. The red arrows indicate the extra β strand found in TgSRS57 and TgSRS16C whereas the black arrow indicates the extended β stands in TgSRS57. Bottom: Left-Superimposition of TgSRS29B dimer (PDB id: 1KZQ) and TgSRS57 showing the first β stand (red arrow) of TgSRS57 that clashes with the other monomer (dark teal) of TgSRS29B dimer, thereby preventing dimerization. Right-Side view of the structural overlay showing the curved β-hairpin loop (dotted box) that impedes access to the potential groove. *Inset*-B*-*factor putty model depicting the mobility of the hairpin loop. **(D)** Top: Top view of the TgSRS29B dimer highlighting the basic groove, Bottom: Same top view of TgSRS57 and TgSRS29B monomer, along the TgSRS29B dimer axis. Electrostatic surface is colored as calculated by APBS [69] (−1 kT/e in red and +1 kT/e in blue); k, Boltzmann’s constant; e, elementary charge.

Comparison of the individual D1 and D2 domains of TgSRS57 to previously characterized members of the SAG1 family, TgSRS29B and TgSRS16C (TgBSR4), revealed that, in contrast to the membrane proximal D2 domains, the membrane distal D1 domains displayed increased structural variability (TgSRS57: TgSRS29B - 2.2 Å rmsd over 130 Cα atoms for D1 and 1.5 Å over 123 Cα atoms for D2; TgSRS57: TgSRS16C – 2.6 Å rmsd over 121 Cα atoms for D1 and 2.0 Å over 135 Cα atoms for D2). Similar to TgSRS16C, TgSRS57 incorporates an additional N-terminal β-strand **(Fig. 4C, indicated by red arrows)** and extended β-strands in the top leaf of D1 compared to TgSRS29B. The two longest strands in the TgSRS57 D1 top leaf are of particular note, as they formed an extended and curved β-hairpin, where the analogous strands in both TgSRS29B and TgSRS16C are much shorter or barely recognized (**Fig. 4C, indicated by black arrows**. These differences are particularly interesting since D1 is apically positioned and thus most likely to engage partners on the host cell surface. Further, the variability in the apical surface may affect dimerization. Homodimerization of TgSRS29B is necessary to form the basic groove that is predicted to coordinate a negatively charged ligand, possibly SPGs [18, 29, 35]. Notably, there was no evidence for dimerization either from the SEC or crystal packing of TgSRS57. In fact, attempts to model the TgSRS57 dimer using the TgSRS29B scaffold were unsuccessful due to a series of steric clashes between the substantially more extensive β sheet structure of TgSRS57 D1 **(Fig. 4C-Bottom left panel)**. Furthermore, the extended and mobile β-hairpin loop of TgSRS57 curved toward the center to guard access to a potential groove **(Fig. 4C-Bottom right panel)**. An electrostatic surface representation of TgSRS57 likewise indicated that it lacks the basic surface groove that, in TgSRS29B, is predicted to coordinate SPGs **(Fig. 4D)**.

### T. gondii heparin-binding proteins are required for optimal infectivity of host cells, but are not necessary for intracellular replication

Previous work demonstrated a dramatically reduced binding of recTgSRS57 to CHO cells pretreated with 30 mM sodium chlorate (NaClO_3_) [26], a potent inhibitor of proteoglycan sulfation [36, 37]. Our current evidence does not support these earlier studies, which concluded that surface coat protein TgSRS57 (along with TgSRS29B) binds cellular SPGs to promote parasite attachment and host cell infectivity. [25, 26, 28]. To better resolve this discrepancy, we investigated whether *T. gondii* parasites bind SPGs in a TgSRS57-dependent manner to promote entry into host cells. RH and *Δsrs57* parasite clones were allowed to invade HFF cells either untreated, or treated with HClO_3_ (to reduce levels of surface associated SPGs). Confluent monolayers of CHO cells, seeded onto 24 well plates, were pretreated with 60 mM of NaClO_3_ for 48 hours, before infecting with 2 × 10^5^ tachyzoites. Parasites were allowed to attach and invade over 30 minutes, washed and left to grow. After 16 hours, the number of vacuoles and total number of intravacuolar parasites was determined across 10 randomly selected fields. In agreement with previous reports [25], the inhibition of proteoglycan sulfation significantly reduced parasite entry to the host cell, supporting a role for SPGs in host cell entry. Compared with non-treated cells, a three-fold reduction in the number of vacuoles was observed in cells pretreated with NaClO_3_, but there was no detectable difference between wild type or Δ*srs57* parasites, suggesting that TgSRS57 does not play a critical role in SPG-dependent attachment or invasion **(Fig. 5A)**. Furthermore, neither the number of vacuoles nor the intracellular replication rate (measured by the average number of parasites/vacuole) was altered in either of the tested strains **(Fig. 5B, 5C)**.

**Figure 5.**
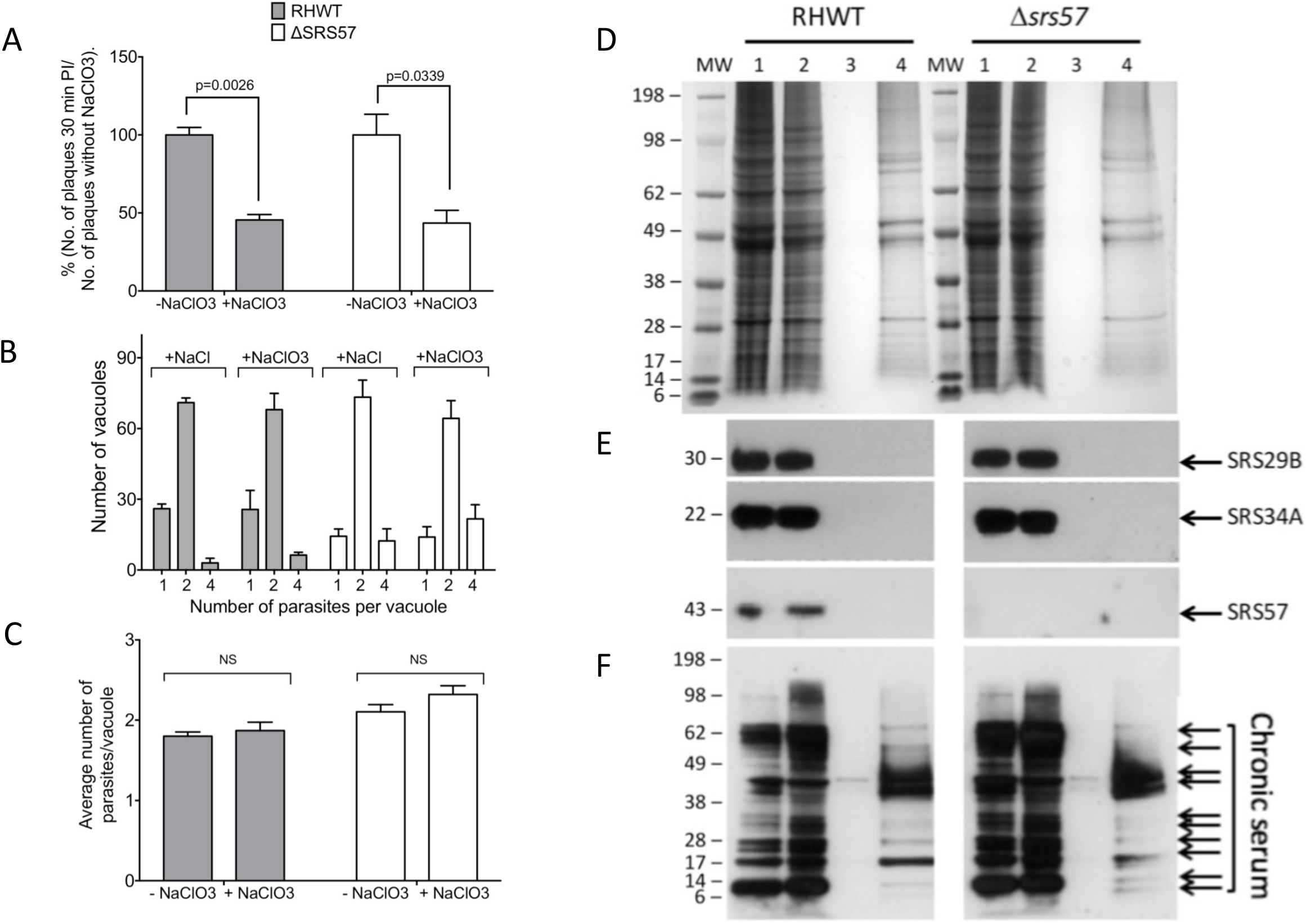
Inhibition of proteoglycan sulfation affects attachment/invasion of T. gondii and the identification of T. gondii heparin-binding proteins. **(A)** 10^3^ tachyzoites of each strain (in triplicate) were inoculated onto HFF cell, pretreated during 48 hrs with 60 mM NaClO_3_ or 60 mM NaCl, and left to invade for 30 min before being washed extensively. After 6 days in culture, the total number of plaques were counted. The same degree of reduction (∼three fold) in plating efficiency was observed in both, RH WT and TgSRS57 deficient strains, demonstrating that, although the infectivity of *T. gondii* is dramatically reduced when the sulfation of GAGs was inhibited, the parasite did not rely on the specific binding of TgSRS57 to SPGs to attach to and invade host cells. The data are expressed as a mean of triplicates ± SD. The statistical significance was assessed by t-test, using Prism 7 software. **(B and C)** Intracellular replication is not compromised by NaClO_3_ inhibition. 2 × 10^5^ tachyzoites of each strain (in triplicate) were inoculated onto HFF cell, pre-treated as in **(A)**. After 12, 18 and 24 hrs. the number of vacuoles containing 1, 2, 4, 8 or 16 parasites/vacuole were counted. No significant differences were observed in the rate of growth of either RH WT or TgSRS57 deficient parasites, when GAG sulfation was inhibited, indicating that intracellular replication is not affected by the inhibition of sulfation. The data are reported as the number of parasites observed per vacuole for 100 vacuoles **(B)** or the average number of parasites per vacuole **(C)** from 3 replicates ± SD. The statistical differences were determined by t-test using Prism 7. **(D)** Pellets of tachyzoites from wild type and knockout strains were lysed in 1% NP-40. Insoluble material was pelleted by centrifugation and the supernatant was subject to heparin-agarose affinity chromatography. Different fractions were analyzed by SDS-PAGE and Western blot. Coomassie blue stained SDS-PAGE showed the protein profile of fractions eluted by heparin-agarose affinity chromatography. Lane 1, loading sample. Lane 2, unbound flow through fraction. Lane 3, wash of the heparin column. Lane 4, high NaCl eluate from the affinity column. Both eluates (WT and knockout) show identical protein profiles. **(E)** Immunoblot of the samples in **(D)** probed with anti-TgSRS29B (top), anti-TgSRS34A (middle) and anti-TgSRS57 (bottom). None of the three SRS proteins bound specifically to heparin. **(F)** Immunoblot of the samples in **(D)** developed with chronic serum from mice infected with *T. gondii* identified at least 10 proteins that bound specifically to the heparin column.

### Identification of T. gondii heparin-binding proteins

We next tested whether native TgSRS57, isolated from the parasite surface, is capable of binding SPGs. This approach allowed us to assay for other *Toxoplasma* surface coat proteins that may specifically bind SPGs. Parasite lysates of RH and the *Δsrs57* strains were subjected to heparin sulfate-agarose affinity chromatography. Fractions collected during the purification process were separated by SDS-PAGE and stained with Coomassie blue (**Fig. 5D**). Approximately 10 proteins were resolved by Coomassie staining of the eluate fraction (lane 4). The eluate fractions were subjected to immunoblot analysis using anti-TgSRS29B (DG52), anti-TgSRS34A (5A6) and anti-TgSRS57 (1F12) monoclonal antibodies. None of these antibodies recognized proteins in the eluate fraction under the conditions utilized (**Fig. 5E**), indicating that these three SRS proteins do not bind sulfated heparin with strong affinity. Probing the same fractions with chronic immune serum from mice infected with the Type II Me49 strain of *T. gondii* established that the 10 bands identified by Coomassie staining, which did not vary between RH and *Δsrs57* strains, were immunogenic, parasite-derived, and possessed a strong affinity for sulfated heparin (**Fig. 5F**). Further studies will be necessary to identify these proteins.

### RHΔsrs57 infected mice show earlier cachexia and increased parasite burden, but no significant difference in survival

Previous work suggested that TgSRS57 was required for infection competency in mice [28]. To assess whether TgSRS57 is a virulence factor during acute infection *in vivo*, three groups of CD-1 mice were challenged intraperitoneally (50 tachyzoites/mouse) with RH, Δ*srs57* and complemented strains respectively. Survival and weight changes were monitored daily. Although there was no significant difference in survival among these groups, an unexpected early presentation of cachexia driven weight loss was detected by day 7 post infection in mice infected with Δ*srs57* parasites compared with the other two groups, in which cachexia driven weight loss started at day 8 **(Fig. 6A)**. This was in contrast to previous studies in which the Δ*srs57* strain CL55 failed to expand *in vivo* or cause weight loss. Strains used in those studies, fRH (French RH) and the mutant CL55 strain, were obtained, cultured and tested in mice using the same conditions used for RH and the Δ*srs57* used herein. Both strains (fRH and CL55) were acutely virulent (**Supplemental Fig. 3)**. In agreement with this observation, bioluminescence imaging during the course of infection revealed a higher parasite burden beginning at day 6 post-infection in mice infected with Δ*srs57* parasites **(Fig. 6B)**. Quantification of the bioluminescence activity confirmed that the increase in parasite load was statistically significant (**Fig. 6C)**. Intriguingly, these differences resolved during the following days and no significant difference in survival was observed **(Fig. 6D)**.

**Figure 6.**
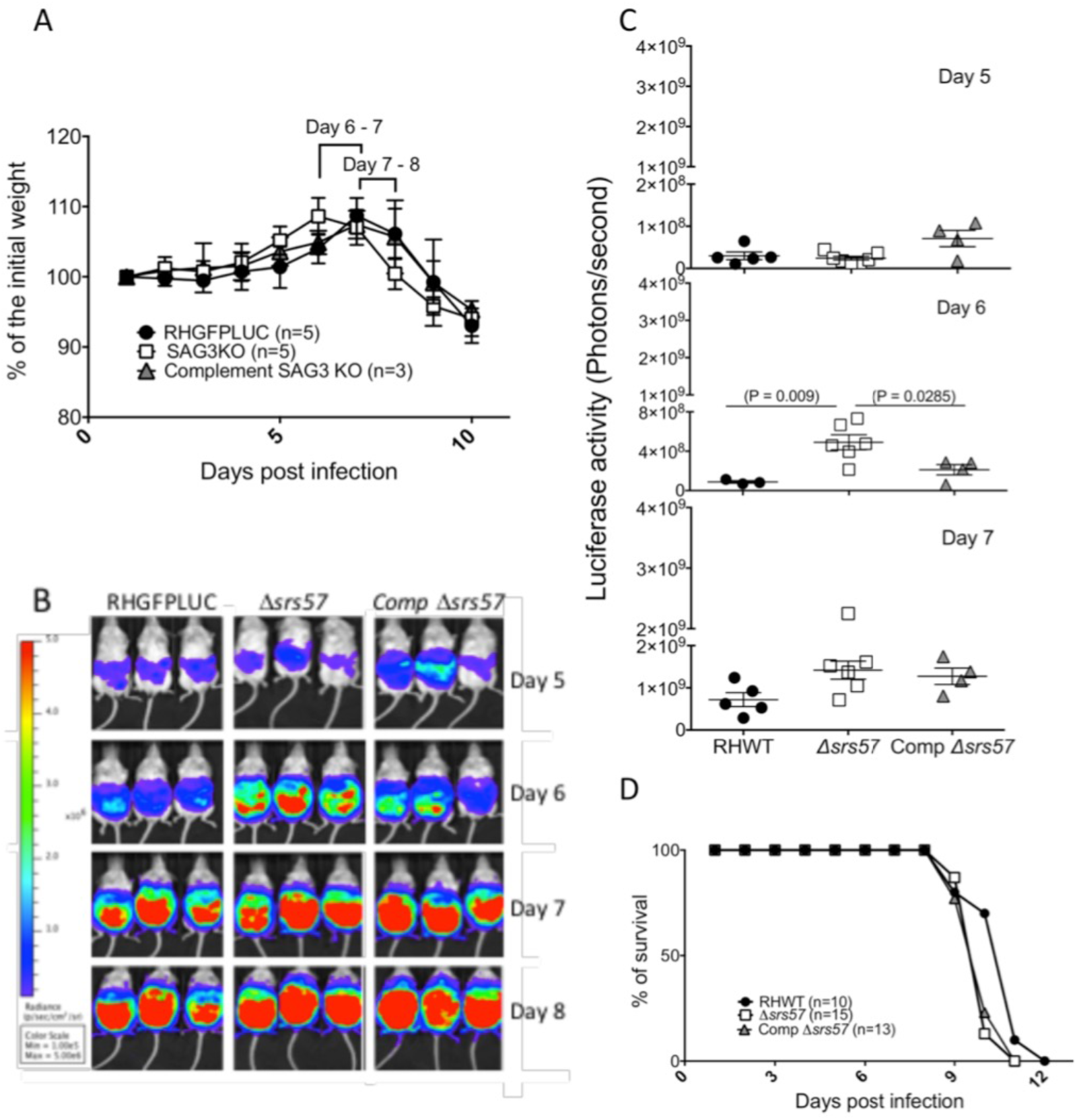
In vivo *characterization of* Δsrs57 *parasites*. **(A)** Loss of weight after murine infection. For each strain, groups of 5, 4 or 3 CD-1 mice were intraperitoneally infected with a dose of 50 parasites/mouse and the daily weight was registered until death. The results correspond to one of three independent experiments. Earlier cachexia was observed in Δ*srs57* parasites, starting at day 6 post infection, compared with RH WT and complemented strains, starting at day 7 post infection. The curves represent the mean ± SD of 5, 5 and 3 animals infected with RH WT, Δ*srs57* and complement strains respectively. **(B)** Bioluminescent detection of parasite burden *in vivo*. CD-1 mice were infected with 50 luciferase-positive WT, Δ*srs57* or complemented strains and imaged daily. Representative images of 3 animals per group, with photon output intensity in photons/second/cm^2^/surface radiance are shown for days 5, 6, 7, and 8. The image shows higher luciferase intensity at day 6 in mice infected with Δ*srs57* parasites, compared with the WT and complemented strains. **(C)** Quantification of the luciferase activity showed differences in parasite burden observed at day 6 post infection in the Δ*srs57* parasites, which was statistically significant (p<0.05) and confirmed the visual increases in luciferase activity observed in **(B)** at day 6, consistent with the loss of weight as shown in **(A). (D)** Survival curve of CD-1 mice challenged with 50 tachyzoites/mouse of the indicated strains. No significant differences were observed between the 3 strains. The results are an aggregate of three independent experiments.

### Type II Δsrs57 parasites are more virulent and exhibit higher parasite burdens in vivo

We next tested whether the absence of TgSRS57 in the CZ1Δ*srs57* strain altered the virulence and parasite expansion kinetics in the Type II genetic background. C57BL/6J female mice were infected with 2×10^3^ parasites intraperitoneally. Survival and weights were monitored daily. Mice infected with CZ1*Δsrs57* parasites died significantly (p=0.014) sooner than those infected with WT parasites (**Fig. 7A**) and exhibited a more pronounced and earlier cachectic response (**Fig. 7B**). These data confirmed the earlier cachexia observed in mice infected with Type I RH*Δsrs57* parasites and demonstrated that SRS57 expression was protective *in vivo*.

**Figure 7.**
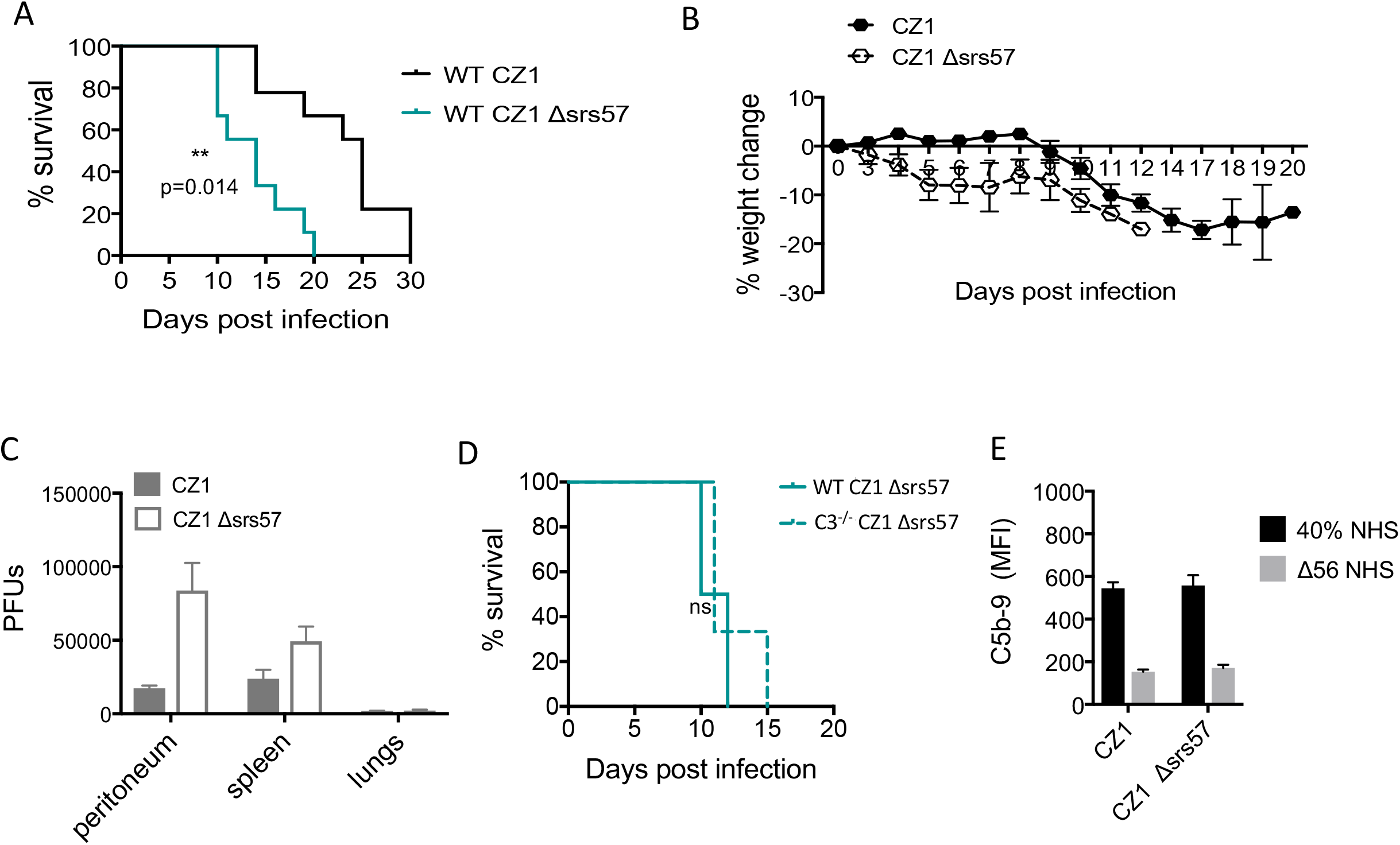
*Mice infected with Type II* Δsrs57 *parasites exhibit altered pathogenesis that is dependent on complement*. **(A)** Survival curve of C57BL/6J mice infected with 2×10^3^ tachyzoites intraperitoneally. Mice infected with *Δsrs57* parasites (teal line) died significantly sooner than those infected with wild type parasites (black line) (p = 0.014). The results are an aggregate of two independent experiments, n=9 per group. **(B)** Weight loss after infection with 2×10^3^ tachyzoites intraperitoneally. Earlier cachexia was observed in *Δsrs57* parasites, starting at day 3 post infection, n=9. **(C)** Parasite burden was determined at day 6 post infection by plaquing peritoneal fluid and homogenized spleens and lungs on HFF cells in 12 well plates. Parasite burden is represented as plaque forming units (PFUs) per organ. Error bars represent mean ± SEM. Data is representative of 2 independent experiments, n=4. **(D)** Survival curve of wild type C57BL/6J and C3^-/-^ mice infected with 2×10^3^ CZ1 Δ*srs57* tachyzoites intraperitoneally. No significant difference (ns) was observed between the two strains of mice. The results are representative of one experiment, n=3 C3^-/-^, n=4 WT. **(E)** CZ1 and Δ*srs57* parasites were incubated in 40% NHS for 30 minutes at 37°C to measure membrane attack complex (MAC) formation by flow cytometry using a monoclonal mouse α-human C5b-9 (1:500, Santa Cruz Biotechnologies).

To determine if parasite burden at day 6 post infection phenocopied the higher parasite burdens observed in mice infected with the RH*Δsrs57* strain, peritoneal fluid, homogenized spleen and lungs were plated onto confluent HFF cells in 12 well plates and plaques were counted as a measure of parasite burden. Mice infected with Type II *Δsrs57* parasites likewise demonstrated significantly higher numbers of parasites in all tissues assayed, demonstrating greater dissemination and parasite burden compared to mice infected with WT parasites (**Fig. 7C**). These data suggest that host immunity induced by the expression of TgSRS57 controls parasite expansion and regulates murine pathogenesis.

### TgSRS57 mediated-protection in vivo is dependent on complement but is not mediated by serum killing of parasites

C3 and its catabolites are critical components of the innate immune system that limit infection by 1) promoting direct lysis of pathogens, 2) targeting them for clearance by phagocytosis, and 3) activating effector arms of the adaptive immune response to control parasite expansion. Our previous study showed that mice deficient in C3 failed to regulate parasite proliferation and were more susceptible to CZ1 infection than WT mice, demonstrating that complement was protective *in vivo* [31]. To test whether the enhanced virulence of CZ1 *Δsrs57* (**Fig. 7A**) was dependent on the presence of C3, WT and C3^-/-^ mice were infected with the CZ1 strain deficient in SRS57 expression, which fails to bind WT levels of C3b and FH. No protective effect was observed in mice replete with C3, suggesting that the interaction between TgSRS57 and C3 is mediating protection *in vivo* **(Fig. 7D)**. Our data support the hypothesis that Type II strains deficient in the surface coat protein TgSRS57 fail to be opsonized with sufficient levels of inactivated C3 necessary to promote a protective response in mice.

Complement-mediated membrane attack complex (MAC) formation followed by pathogen lysis is one mechanism of protection that directly controls pathogen proliferation. To test whether parasite resistance to serum killing was dependent on TgSRS57 expression, *Δsrs57* parasites were incubated in 40% NHS for 30 minutes, and parasite viability and membrane attack complex formation (C5b-9) were measured by flow cytometry. *Δsrs57* and WT parasites were equally resistant to human serum killing (data not shown) and had similar levels of C5b-9 MAC formation (**Fig. 7E**). These data do not support complement mediated lysis as the major complement effector function controlling serum resistance, *T. gondii* infection load, or acute virulence in the mouse model. Hence, opsonization of TgSRS57 appears to be a critical step in the host protective phenotype *in vivo*. These data also suggest that C3 split product effectors and other soluble factors associated with the complement pathway are more critical to promote this protective phenotype.

### TgSRS57 regulates systemic and local pro-inflammatory cytokine responses

Several studies have shown that Type I parasites exhibit greater dissemination and parasite burdens compared to Type II strains *in vivo*, and induce a dysregulated Th1 inflammatory cytokine production that significantly contributes to mortality [38, 39]. To investigate whether inflammatory cytokine production was a factor contributing to the altered virulence kinetics, C57BL6/J mice infected with 2×10^3^ Type II *Δsrs57* parasites intraperitoneally were sacrificed 6 days post infection. Serum and splenocytes were harvested and used to measure IL-12, IL-6, TNF-α, IFN-γ cytokine production. Proinflammatory cytokines in the serum **(Fig 8A)** and those produced locally by *ex vivo* splenocyte cultures **(Fig 8B)** were significantly higher in the absence of SRS57. This dysregulated cytokine response is likely explained by the elevated parasite burden in the peritoneum, spleen and lungs compared to wild type mice. However, additional studies are required to determine if the increased virulence of *Δsrs57* parasites is a necrotic process driven by the failure to control parasite proliferation, or is the result of the differential induction of the cytokine response by the absence of TgSRS57. In effect, the expression of TgSRS57 is a resistance factor that promotes disease tolerance, host survival, parasite persistence and transmissibility.

**Figure 8.**
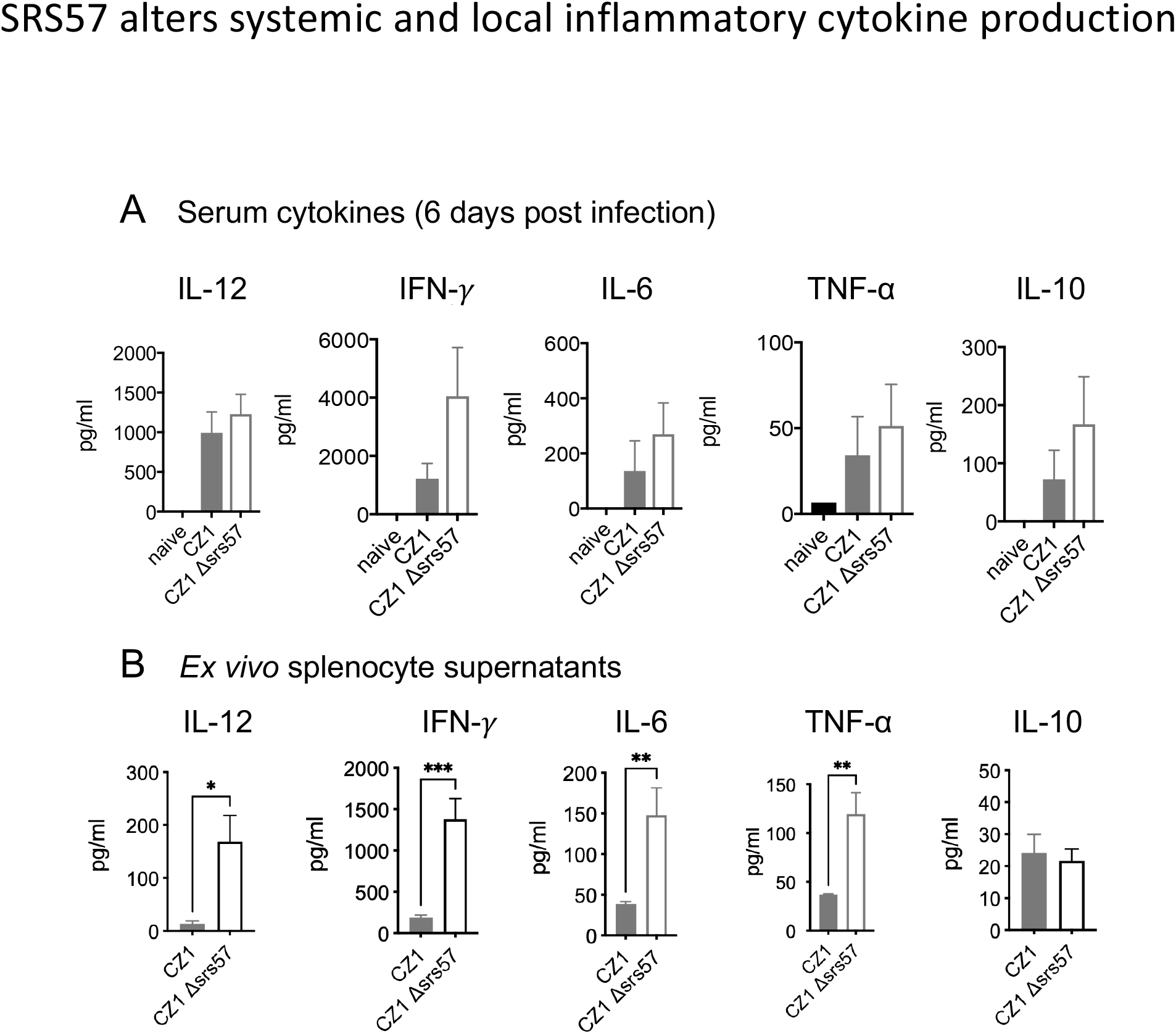
TgSRS57 alters systemic and local inflammatory cytokine production. 6-8-week-old C57BL6 mice were infected with 2×10^3^ CZ1 and Δ*srs57* tachyzoites intraperitoneally and sacrificed 6 days post infection. (A) Sera and (B) supernatants from *ex vivo* splenocytes that were cultured for 24 were collected at 6 days post infection for measuring IL-12p40, IFN-γ, IL-6, TNF-α, and IL-10 by ELISA (R&D Systems). Data is representative of 2 independent experiments, n=4. Error bars represent mean ± SEM and p-values were obtained using a Student t-test, *p<0.05, **p<0.01, ***p<0.001.

## DISCUSSION

Host factors, including components of innate immunity such as IgM [40], complement proteins [30, 31], and host sulfated proteoglycans [26], have been shown to interact with the surface coat of *T. gondii*. Several studies previously implicated SRS proteins in these interactions, but their contribution to *T. gondii* pathogenesis were unclear. This study investigated the specific interaction of surface protein TgSRS57 with SPGs, C3b and FH in order to gain a better understanding of its biological function. We showed that expression of TgSRS57 regulates the recruitment of FH, the deposition of the complement effector protein C3b, and that this interaction specifically plays an important role in the pathogenesis of *T. gondii* infection in mice. The reduced capacity of TgSRS57 to interact with FH, however, was not explained by a specific interaction with SPGs, and we provided a structural blueprint to support this conclusion. TgSRS57 is not required for host cell attachment, the productive invasion of host cells, nor does it possess a lectin-specific activity for cell surface associated glycosaminoglycans, as previously suggested [26, 28]. We did, however, identify approximately 10 parasite proteins that specifically react with SPGs, and that SPGs are required for optimal parasite invasion and growth. Thus, it is possible that TgSRS57 interacts with other parasite proteins that bind SPGs to coordinate or present FH in a way that facilitates C3b inactivation. Notably, *in vivo* characterization of the *Δsrs57* parasites established that interactions between TgSRS57 and C3 play a critical role in in promoting disease tolerance by regulating parasite virulence and inducing parasite persistence.

Intracellular pathogens have evolved multiple strategies to gain entry to their target host cells. Whether the invasion process is passive or active, the initial recognition and attachment phase is critical for the success of internalization. A variety of cell surface effector proteins from both invading organisms and their hosts participate in this process. Many viruses, bacteria, fungi and parasites express lectins that bind to cell surface associated SPGs such as heparan sulfate to promote initial recognition, attachment and cell entry [8]. It is also known that microbial pathogens utilize these interactions to evade and subvert host defense mechanisms, to their own advantage, in order to survive within the host [41]. SPGs are expressed in virtually all cell types, from simple invertebrates to humans [42]. They are glycoproteins that contain one or more covalently attached linear polysaccharide chain and are referred to as glycosaminoglycans (GAGs). The two major types of GAGs found in animal cells are heparan and chondroitin sulfate [8]. Chondroitin sulfate A (CSA) is a cell surface receptor for *Plasmodium falciparum*-infected erythrocytes [43] and cytoadhesion of parasitized erythrocytes to the placenta is mediated by the specific interaction of a parasite encoded adhesin VAR2CSA with CSA present in placental proteoglycans [44]. *Plasmodium berghei* uses the sulfation level of heparan sulfate proteoglycans (HSPGs) to navigate within the mammalian host. High levels of these glycoproteins on the surface of hepatocytes activate invasion, whereas low levels promote migration from the skin to the liver [45]. The transition from a migratory to an invasive phenotype is mediated by a signaling pathway, which includes a calcium dependent protein kinase 6. Binding of the sporozoite’s major surface protein, the circumsporozite protein (CSP) triggers this process [45].

The first evidence that *T. gondii* possesses a sugar binding activity came from an early investigation in which tachyzoites were shown to bind the neoglycoprotein bovine serum albumin-glucosamide [46]. It was later demonstrated that attachment of the parasite to its target cell is mediated by parasite lectins that bind GAGs [11]. At least three surface adhesins that resolved by SDS-PAGE at 26, 45 and 65 kDa were identified for their lectin-specific activity and by their ability to bind to the surface of erythrocytes and HFF cells, although the molecular and biochemical characterization of these lectins, and their putative target sugars remain undefined [11].

Despite the fact that the active process of internalization is conserved throughout the phylum, notable differences in the spectrum of invaded host cells is observed between different members of the phylum. *Plasmodium spp*. infect host hepatocytes and erythrocytes, whereas *Cryptosporidium spp*. infect only enterocytes and *Toxoplasma* possesses the unique characteristic of infecting virtually any nucleated cell [15, 17]. It is tempting to speculate that a correlation between host range and the number and variability of surface adhesins exists. For example, more than 180 surface-associated SRS adhesin-encoding genes have been identified in the genome of *Toxoplasma* [16], whereas *Plasmodium* only possesses 14 members of the related 6-Cys protein family that, like *Toxoplasma* SRS proteins, are expressed developmentally, with different sets of 6-Cys proteins expressed at different stages throughout the parasite life cycle. Both families, the 6-Cys and SRS proteins are derived from a common ancestor within the Apicomplexa and each have evolved their own adhesion superfamily of surface coat antigens based on distinct structural domains referred to as the SRS and s48/45 domains in *Toxoplasma* and *Plasmodium*, respectively [20]. While relatively little is known about the specificity of the SRS proteins, three members of the analogous 6-Cys family play a role in gamete fertility (p230, p230p and p47) [47] although it is not clear whether these fertility factors interact via lectin binding. Other 6-Cys proteins are known to facilitate invasion of hepatocytes (P52/P36) and promote binding to CD36 (Sequestrin) and Factor H (Pf92) [20, 21, 23, 24].

In addition to attachment and adhesion, evasion of host immunity is critical to establishing a successful infection. Previously, we showed that the surface coat of *T. gondii* regulates serum resistance by recruiting host proteins Factor H and C4b-binding protein to downregulate alternative and lectin pathways of the complement system, respectively [31]. Serum resistance was predominantly attributed to the regulation of the alternative pathway by Factor H [31]. Factor H is composed of 20 homologous complement control protein (CCP) domains, otherwise known as short consensus repeats, that give the appearance of flexible “beads on a string” that can fold back on itself [48]. Factor H predominantly interacts with C3b and host molecules such as sialic acid and SPGs to discriminate between self and non-self surfaces. With several binding sites for proteoglycans and C3b, structural studies have established that Factor H simultaneously binds negatively charged anions like SPGs and the C3d portion of the C3b molecule [34] on host surfaces to mediate protection against complement. Although this study established that TgSRS57 regulates C3b opsonization at the parasite surface, and that, in its absence, recruitment of FH to the parasite surface is reduced, it does not provide precise mechanistic insight for FH recruitment, suggesting that other proteins may be required for this interaction. It is also not clear how TgSRS57 is recognized by proteins of the lectin pathway, such as the mannose binding lectin, to facilitate C3b deposition. TgSRS57 lacks predicted N-glycosylation sites, and is not thought to be modified by other glycans, however O-linked glycosylation cannot be ruled out. It is possible that TgSRS57 heterodimerizes with another SRS protein that possesses lectin activity, similar to the structurally related 6-CYS proteins from *Plasmodium* that heterodimerize, but thus far, no evidence exists for SRS heterodimerization.

Previous studies suggested that an array of *T. gondii* surface and secreted proteins, including TgSRS57, TgSRS29B, ROP2, ROP4, and GRA2 bind heparin [25, 26], a pharmaceutical mimic of heparan sulfate (HS), that is a highly-sulfated glycosaminoglycan, found *in vivo* in mast cells, and within connective tissue. It differs from HS in the degree of its sulfation, with heparin displaying higher N and O-sulfation than HS [49] and it is often used to study interactions between pathogens and host cells. A homology model of TgSRS57 based on the structure of TgSRS29B suggested a possible SPG-binding ability for TgSRS57 through a positively charged binding cleft. In order to validate this model, and to investigate the structural and molecular basis of TgSRS57-SPG interaction, we determined the high-resolution crystal structure of TgSRS57. We found that TgSRS57 shared the same overall architectural features with previously characterized SRS proteins. However, and in contrast to the homology model, our structural data did not identify a positively charged binding cleft capable for accommodating SPGs. Moreover, our binding studies with native or recombinantly expressed TgSRS57 failed to indicate any binding with heparin. It is interesting to note that binding studies in the past used recombinantly expressed His-tagged proteins to show TgSRS57-SPG interactions (23, 24) but the presence of the 6×His may non-specifically bind to HS, [50], giving rise to false positive results. In our study herein, the TgSRS57 deficient parasites did show a reduced capacity to recruit FH, which is thought to be recruited to host cell surfaces by binding SPGs, so additional studies are required to identify the precise molecular details that promote TgSRS57-dependent recruitment and binding of FH to the parasite surface coat.

Because the functional relevance of TgSRS57 remained elusive, we further characterized SRS57 deficient strains *in vitro* and *in vivo*. The TgSRS57 deficient strain generated in the Type I RH background, and the previously published CL55 Δ*srs57* mutant, both failed to identify any difference in extracellular survival, adhesion, intracellular replication or a change in murine virulence. This is in stark contrast to the previously published results for this knockout, which identified an adhesive role for TgSRS57 that facilitated the invasion process and virulence in mice [26, 28]. Although we did observe a difference in dissemination kinetics *in vivo* for the TgSRS57 deficient Type I strains, this did not result in a dramatic change in the kinetics or time to death. While it is formally possible that the differences observed in host susceptibility between the two studies are the result of the type of mouse strain used, as it has been shown previously that mouse genetics can influence both acute mortality and cyst load [51], the fact that our infections used the more resistant CD-1 outbred strain, whereas the previous work was done using susceptible inbred Balb/C lines, argues against this as a likely explanation.

We also generated a *Δsrs57* strain in the less virulent Type II CZ1 strain and infected susceptible C57BL/6J mice to further investigate differences in parasite burden and cachexia as was observed during infections with the Type I deficient strain. The CZ1*Δsrs57* parasites were significantly more virulent, which was opposite to the protective phenotype previously described [28]. Infected mice exhibited higher parasite burdens that promoted a more vigorous Th1 pro-inflammatory response, suggesting that the ability of TgSRS57 to regulate C3b deposition and itw inactivation attenuates virulence, presumably by regulating parasite proliferation during acute infection. However, it was not clear the extent to which uncontrolled parasite expansion versus immunopathology contributed to the rapid mortality phenotype observed in mice infected with Δ *srs57* parasites. IgM, an important factor of humoral immunity, was previously shown to limit parasite proliferation by preventing host cell invasion (37), which established a protective role for humoral immunity. Separately, opsonized parasites are known to enter host cells by a phagocytic process and they retain replication competence [52, 53], which has been postulated to induce host tolerance to acute mortality by virulent strains of *T. gondii* [53].

Because TgSRS57 regulates C3b deposition on the parasite surface, the biological relevance of C3 during acute infection was evaluated. C3^-/-^ mice infected with TgSRS57 sufficient WT parasites were more susceptible, indicating that complement was protective during acute infection. However, no difference in virulence was observed between wild type mice and C3^-/-^ infected with Δ *srs57* parasites, confirming that the interaction between TgSRS57 and C3 are needed for disease tolerance and host protection *in vivo*. Because we have no evidence that TgSRS57 is covalently modified by C3, our data suggest that a factor independent of TgSRS57 regulates the level and inactivation state of C3 on the parasite surface to promote the host protective response in mice. The complement system is known to cross talk with several other innate recognition systems, including TLR receptor signaling [54, 55] and C-type lectin receptor Dectin-1 [56] to coordinate immunological responses [57], so it is possible that TgSRS57 might coactivate complement and one of these systems to regulate virulence and control parasite proliferation, as has been observed previously for TgMIC1 [13].

Since TgSRS57 expression did not impact MAC complex formation nor complemented-mediated lysis, it is likely that additional complement effector functions, such as complement-mediated phagocytosis or stimulation of adaptive immunity, was affected by the reduction of C3b deposition on *Δsrs57* parasites, or in the case of C3 deficiency. Importantly, the loss of complement mediated protection in these mice occurred during the induction of adaptive immunity (before day 21), indicating that complement likely plays a contributing role in the promotion and regulation of adaptive immune responses. For example, C3 split product effectors C3b and anaphylatoxin C3a play important roles regulating B and T cell responses, respectively. C3a has recently been shown to play a critical role regulating Th1 responses [58, 59], which are essential for resolving acute *T. gondii* infection. In addition, antigens covalently bound by catabolites iC3b and C3dg, are recognized by CR2 on B cells, lowering the threshold for B cell activation and promoting antibody production [60, 61]. In the absence of TgSRS57 or complement, it is possible that infected mice exhibited an altered TgSRS57-specific B cell response which resulted in less effective B cell immunity. Further studies are required to determine which C3 downstream effector proteins regulate the enhanced mortality observed in C3 deficient animals and what role TgSRS57 plays in such immune modulation.

In aggregate, our data indicate that TgSRS57 does not possess a lectin-like activity nor does it directly mediate attachment through an SPG host cell receptor interaction. However, approximately 10 *T. gondii* lectins were identified that specifically interact with heparin. Ongoing studies to identify the nature of these proteins will shed light into the mechanisms of initial host cell recognition and the possible mechanisms of TgSRS57-dependent FH recruitment. *In vitro* and *in vivo* studies revealed a novel function for TgSRS57 through its interaction with C3b to promote disease tolerance by regulating both parasite virulence and persistence in the mouse model.

## MATERIALS AND METHODS

### Parasite strains, host cells and cell culture conditions

Wild type RH tachyzoites, knockout and complement strains were propagated in human foreskin fibroblasts (HFF) in Dulbecco’s Modified Eagle Medium (DMEM) supplemented with 10% heat inactivated fetal bovine serum (FBS), 2 mM glutamine, 10 μg/ml of gentamicin and 0.5% of penicillin/streptomycin (100X) and maintained at 37°C in a humid 5% CO_2_ atmosphere.

### Targeted disruption and complementation of the TgSRS57 locus

The targeting vector used to generate the Δ*srs57* strain was derived from the pminiHXGPRT vector (Catalogue Number: 2855. NIH AIDS Reagent program, Contributor: David Roos). The 5’ flanking (flk) sequence (2.0 Kb) and 3’ flk sequence (3.1 Kb) were amplified from RH genomic DNA and cloned into KpnI/HindII and BamH1/SacI sites, respectively. The primers to amplify the 5-flanking sequence were 5SRS57FW (GCGGTACCAAGTCGAAGA GTGCGTTCGT) and 5SRS57Rev (GCAAGC TTAAGGATACCGTGTGCGAAAAC) and the primers to amplify the 3’-flanking sequence were 3SRS57FW (GCGGTACCGT GTGAGACCCTCTCT) and 3SRS57Rev (GC AAGCTTACGAATCCAGTGCAGTTTCC).

The 5’dhfr-HXGPRT-3’dhfr cassette (1.9 Kb) contained in the vector was used as a marker for positive selection. The 5’flk TgSRS57/dhfr-HXGPRT-dhfr/3’flk TgSRS57 construct (7.0 Kb) was released from the vector using the restriction enzymes KpnI and SacI. 10^7^ RHΔHXGPRT parasites tachyzoites were transfected with 50 μg of the construct. Transfections were performed by resuspending parasites and DNA in electroporation buffer (120 mM KCl, 0.15 mM CaCl_2_, 10 mM K2HPO_4_/KH2PO_4_. pH 7.6, 25 mM HEPES, pH 7.6, 2 mM EDTA, 5 mM MgCl_2_, 2 mM ATP, 5 mM glutathione), using a BTX model ECM630. 24 hrs after transfection parasites were selected with 50 μg/ml mycophenolic acid and 50 μg/ml xanthine. Surviving populations were cloned by limiting dilution and screened for the loss of the TgSRS57 expression by flow cytometry. For complementation, 10^7^ parasites knockout were co-transfected with 50 μg of the TOPO 4.0 vector harboring a PCR amplified TgSRS57 genomic locus including the promoter and the 5’ and 3’ untranslated regions (3.7 Kb) plus 5 μg of a pGFP-LUC plasmid, containing cDNA sequences of the Green Fluorescence Protein (GFP) and Fire Fly Luciferase (LUC). The primers used for the amplification of the TgSRS57 locus were PF: CAAGACACACAAGTGCGTGTGATG AC and PR: CACTGTTCTGATGGAATGCT CGCTAC and TA cloned into the TOPO 4.0 vector. Before transfection both plasmids were linearized with Not1. 10 days post-transfection, stable parasites expressing GFP were FACS sorted onto 96 well plates. Positive clones expressing GFP were analyzed for TgSRS57 surface expression by flow cytometry.

### Generation of CZ1 Δsrs57 mutant strains using CRISPR/Cas system

A clonal isolate of CZ1-Δ*ku80*/Δ*hpt* was used to generate the CZ1-Δ*ku80*/Δ*srs57* parasites. The *TgSrs57* gene was replaced by the drug selectable marker HPT (*HXGPRT* – hypoxanthine-xanthine-guanine phosphoribosyl transferase) flanked by 40bp homologies to *TgSRS57* using CRISPR/Cas system (*SRS57* guide RNA sequence: GTGTGTGTTGTCTGCGATCT). Fifteen micrograms of *HXGPRT* repair oligo and 30 μg of guide RNA were transfected into by electroporation using the BTX system. The parasites were then allowed to infect T25 flask of HFFs for overnight, after which the medium was changed to mycophenolic acid (MPA) and xanthine at 50ug/ml for HXGPRT positive selection. After two passages, the parasites were cloned using single-cell method into 96 well plates. Δ*srs57* clones were identified by PCR of the integrated of *HXGPRT* into the open reading frame using the following primers: (*TgSRS57* forward primer: CTTTTTCAGTAACGCTTCGT and HXGPRT reverse primer: GAGCACCCTGATATGACAAT). TgSRS57 deletion was further confirmed by staining and analyzed by flow cytometry and by western blot.

### Flow cytometry

Suspensions of freshly harvested tachyzoites from the different strains were washed with PBS, fixed for 15 min with 2% (w/v) formaldehyde in PBS at room temperature (RT) and stained with the primary monoclonal antibody 1F12 (specific for the TgSRS57 protein), diluted 1:4000, washed three times with FACS buffer (1% BSA in PBS) and stained with the secondary antibody anti-mouse conjugated with Allophycocyanin (APC), diluted 1:200. After 3 washes in FACS buffer, the stained parasites were analyzed for surface protein expression by flow cytometry. Data were collected using a FACS Canto flow cytometer (BD Bioscience) and analyzed using FlowJo software.

### SDS-PAGE electrophoresis and Western blot

Lysate, flow through, last wash and eluate fractions were analyzed by SDS-PAGE on Novex bis-tris 4 to 12% gradient gels (Invitrogen) using MOPS SDS as running buffer. Samples were heated at 70°C for 10 min in sample buffer containing 50mM dithiothreitol as reducing agent. Gels were stained with Coomassie Blue. For Western blot, the gels were electro transferred to nitrocellulose over night at 10V constant, using a Mini Trans-Blot® Electrophoretic Transfer Cell (BioRad). After transfer, the membranes were blocked in PBS containing 5% non-fat dry milk and 0.05% Tween 20 prior the addition of the appropriate antibodies (anti-TgSRS29B (DG52) mAb, 1:5000; anti-TgSRS34A (5A6) mAb, 1:400; anti-TgSRS57 (1F12) mAb, 1:1000 or mouse chronic serum after 5 weeks post infection. After three washings with PBS containing 1% non-fat dry milk and 0.05% Tween 20, the membranes were incubated with peroxidase conjugated goat anti-mouse antibody (Sigma), diluted 1:1000), washed three times and visualized by chemiluminescence. Alternatively, parasites extracts were treated with GPI-specific phospholipase C (GPI-PLC). Briefly, parasites were washed with PBS, resuspended in 50 ml of lysis buffer (50 mM Tris [pH 8.0], 5 mM EDTA, 5% NP-40) and incubated with GPI-PLC from *Bacillus cereus* for 1 h at 37°C. Cross-reacting determinants (CRDs) of GPI anchors, which were unmasked by this treatment, were detected with a polyclonal antiserum against CRD.

### Extracellular survival

Suspensions of free parasites (10^3^ tachyzoites/ml) were incubated in culture media at 37 °C with 5.5% CO_2_, in absence of host cells for 0, 1/2, 1, 2, 4, 8, 24 and 48 hrs prior infection. After the indicated times a total of 10^3^ tachyzoites of each strain was applied to a fresh monolayer of HFF cells, seeded onto 24 well plates and incubated during 5 days in normal culture condition without disturbance. After incubation, the wells were rinsed with PBS, fixed with 100% of methanol for 5 min and stained with crystal violet. Parasites survival was estimated by counting the number of plaques. The assay was carried out in triplicates. The results were presented as the mean ± SD (n=3).

### Plating efficiency

10^3^ freshly lysed tachyzoites were allowed to invade monolayers of HFF cells for 30, 60, and 90 min and 24hrs. After the indicated times, uninvaded parasites were removed by washing four times with PBS and allowed to replicate undisturbed at 37°C and 5.5% CO_2_ atmosphere for 6 days. The total number of plaques formed at each time point were counted. The plating efficiency was expressed as a percentage of the total number of plaques at 24 hrs. The experiment was performed in triplicate and the statistical differences among means were calculated using the One-way ANOVA (non-parametric) test.

### Rate of replication

Confluent monolayers of HFF cells, grown in 12 well plates, were infected with 10^5^ tachyzoites/well. After 4 hrs of incubation at 37°C and 5.5% CO_2_, to allow the parasites to invade, the wells were washed 3 times with PBS and incubated for 16, 24, and 36 hrs in regular media. At each different time the cells were fixed with 100% of methanol for 10 min and the average number of parasites/vacuole was scored by counting 100 randomly selected vacuoles. The results are presented as the mean of two independent experiments ± SD.

### Flow cytometry-based complement assays for C3b deposition, FH recruitment, C5b-9 formation and parasite viability

Complement deposition was measured by flow cytometry as previously described [31]. Briefly, parasites were collected from a freshly lysed monolayer of human foreskin fibroblast cells, filtered, and washed twice with PBS to remove media. 1×10^6^ parasites per time point were incubated with 10% non-immune human serum (pooled NHS, Cederlane) in complement activating buffer HBSS^++^ (Hanks Buffered Saline Solution, 1mM MgCl_2_, 0.15mM CaCl_2_). Complement activation was stopped using cold PBS and washed twice with cold PBS to remove excess serum. For viability assays, parasites were exposed to 40% NHS, washed three times with cold PBS to remove excess serum, and stained with a fixable viability stain (Life Technologies) for 15 minutes and washed twice with PBS. Parasites were then fixed with 1% PFA solution for 10 minutes, washed twice, and stained with mouse α-human C3b/iC3b antibody (1:500) (Cedarlane), mouse α-human C5b-9 (1:500), (Santa Cruz Biotechnology) or goat α -human Factor H (1:2,000) (CompTech), followed by α-mouse APC (eBiosciences) or α-goat APC conjugated antibody (Jackson Laboratories), both at 1:1,000. Antibodies were diluted in FACs buffer (PBS + 1% BSA), incubated for 30 minutes at room temperature and protected from light, followed two washes with FACs buffer. 20,000 events were collected on a BD Fortessa flow cytometer and analyzed by FACs DIVA, FlowJo and Prism software

### Cloning, expression and purification of TgSRS57

The TgSRS57 sequence from the signal peptide cleavage site (SP) to GPI anchor site (GPI) was codon optimized, synthesized by GenScript, and subcloned into a modified pAcGP67b vector (Pharmingen) incorporating a C-terminal hexahistidine tag and thrombin cleavage site. TgSRS57 construct extending from the end of the predicted signal peptide to the end of predicted D2 was cloned using PCR. TgSRS57 clones were transfected with linearized Baculovirus DNA into Sf9 cells and amplified to a high titre. Hi-5 cells at 1.8 × 10^6^ cells ml^-1^ were infected with amplified virus for 65 hrs, after which time the supernatant was harvested, concentrated and applied to a HisTrapFF Ni affinity column. TgSRS57 was eluted with an increasing concentration of imidazole and fractions analyzed by SDS-PAGE and pooled based on purity. The hexahistidine tag was removed by thrombin cleavage and TgSRS57 was further purified by size exclusion chromatography (Superdex 16/60 75) in HEPES buffered saline (HBS – 20 mM HEPES pH 7.5, 150 mM NaCl).

### Crystallization and data collection

Diffraction quality crystals of TgSRS57 were grown in 25% PEG1500. The final drops consisted of 0.6 μl protein (12 mg ml^-1^) with 0.6 μl reservoir solution and were equilibrated against 100 μl of reservoir solution. Cryo protection was carried out in mother liquor supplemented with 25% glycerol for 20 seconds and the crystal was flash cooled at 100 K directly in the cryo stream. Diffraction data were collected on beamline 9-2 at SSRL (Stanford Synchrotron Radiation Laboratory).

### Data processing, structure solution and refinement

Diffraction data to a resolution of 1.59 Å were processed using Imosflm [62] and Scala [63] in the CCP4 suite of programs [64]. Initial phases were obtained by molecular replacement (MR) using PHASER [65] with the individual D1 and D2 domains of TgSRS29B (PDB 1KZQ) pruned with CHAINSAW [66]. Solvent molecules were selected using COOT [67] and refinement carried out using Refmac5 [68]. Stereo-chemical analysis performed with PROCHECK and SFCHECK in CCP4 [64] showed excellent stereochemistry with more than 97% of the residues in the favored conformations and no residues in disallowed orientations of the Ramachandran plot. Overall 5% of the reflections were set aside for calculation of R_free_. Data collection and refinement statistics are presented in Table S1. The atomic coordinate and structure factor files have been deposited in the PDB under accession code 8DDP.

Electrostatic surface images were calculated using Adaptive Poisson-Boltzmann Solver (APBS) electrostatic surface evaluation tool with default parameters (Ionic concentration of 0.150 mol/l NaCl, biomolecular dielectric constant of 2 and solvent dielectric constant of 78.54 (water)). The charges and radii of atoms in the PQR file required by APBS were taken from AMBER94 force field [69].

### Inhibition of Glycosaminoglycan sulfation

HFF cells were seeded in 24 well plates and cultured in DMEM supplemented with 10% FBS and 60 mM NaClO_3_, a reversible inhibitor of adenosine 39-phosphoadeny-lylphosphosulfate and proteoglycan sulfation or 60 mM NaCl as a control. After 48 hrs the monolayers were washed once with PBS. For attachment and invasion assays each well was inoculated with 10^3^ tachyzoites suspended in complete DMEM and the plates were centrifuged for 2 min at 500 g to accelerate the contact of parasites with the host cells. After 1hr incubation at 37°C and 5.5% CO_2_ atmosphere, wells were washed 3 times with PBS and incubated undisturbed, in regular culture conditions. After ∼ 6 days, the total number of plaques in each well were counted. The results were expressed as a mean of triplicates ± SD. Paired t-test, using Prism 7 software, was applied to analyze statistical differences between conditions. For replication assays, wells were inoculated with 2 × 10^5^ tachyzoites, treated as above and incubated for 4 hrs in complete DMEM, supplemented with 30 mM NaClO_3_ or 30 mM NaCl. After 12, 18 and 24 hrs the number of vacuoles containing 1, 2, 4, 8 or 16 parasites/vacuole were counted. The data are reported as the average number of parasites per vacuole from three replicates ± SD. The statistical differences were determined by t-test using Prism 7.

### Heparin binding assay

To isolate heparin binding proteins, 10^8^ freshly harvested tachyzoites were washed with PBS and then lysed for 30 min on ice in 100 μl of lysis buffer (1% of NP-40, 0.25M NaCl in phosphate buffer pH 7.2). The lysates were centrifugated at 14.000 rpm for 30 min. Cleared lysates were applied to a heparin-agarose type I beads (Sigma) previously equilibrated with 0.2% NP-40 in phosphate buffer, pH 7.2 and incubated for 30 minutes under gently rotation. Bound beads were extensively washed with 0.2% NP-40, 0.5M NaCl in phosphate buffer pH 7.2, followed by elution of the heparin bound proteins with a high salt elution buffer (0.2% NP-40, 1M NaCl in phosphate buffer pH 7.2).

### Mice infections

Outbred female CD-1 mice (from Charles River), 6-8 weeks old, were used for all *in vivo* experiments. Groups of 5 mice of each strain were intraperitoneally infected with 50 freshly harvested tachyzoites. The cachexia induced during the course of the acute infection was monitored daily. All procedures were performed in accordance with the guidelines of the National Institutes of Health.

### In vivo imaging

For living image, mice were injected with 150 ul (3 mg) of D-luciferin substrate, maintained at least 10 min to allow dissemination of the substrate and anesthetized in an oxygen-rich induction chamber with 2% isofluorane. Images were collected during 5 min using an IVIS (In Vivo Imaging System) Spectrum (PerkinElmer). Mice were imaged at days 5, 6, 7, and 8 post infection. Data acquisition and analysis was performed using the living imaging software.

### Plaque assays to quantify parasite load

Mice infected with CZ1 strains were euthanized at day 6 post infection. Peritoneal fluid was collected by injecting 5ml of sterile PBS into the peritoneal cavity of euthanized mice. Spleens and lungs were harvested and homogenized over 70-μm strainers in 5ml of X-VIVO-20 serum free media. Ten-fold serial dilutions starting with 500μl of resuspended homogenate were added to HFF monolayers in 12 well plates and cultured undisturbed in DMEM media for 7-9 days. HFFs were fixed in methanol and stained with crystal violet. Plaques were counted using an inverted microscope and parasite loads were expressed as CFU per organ.

### Ex vivo splenocyte cultures

Spleens were homogenized over 70-μm strainers in 5ml of X-VIVO-20 serum free media. After red blood cell lysis (commercial RBC lysis buffer), splenocytes were plated in a 96-well flat bottom plate at 5×10^5^ splenocytes per well in 250μl of X-VIVO serum free media and cultured for 24 hours. 250μl of supernatants were collected for analysis.

### Measuring cytokine production from serum and ex vivo splenocyte by ELISA

After euthanasia, sera was prepared from blood collected by cardiac puncture. Blood was allowed to clot for 30 minutes and then spun at 1000rpm for 15 minutes to extract serum. Supernatants of *ex vivo* splenocyte cultures were collected as described above. Samples were analyzed using ELISA kits for IL-12p40, IFN-γ, TNF-α, IL-6, IL-18 and IL-10 according to manufacturer’s instructions (R&D Systems).

## Acknowledgements

We thank the staff at the Stanford Synchrotron Radiation Laboratory for their expert contributions, and Vanessa Lloyd and Emily Jansen for their initial work generating constructs to disrupt TgSRS57. We are grateful to Dr. Stan Tomavo for providing the French RHΔ*srs57* parasite CL55, and the anti-TgSRS57 monoclonal antibody (1F12). This work was financially supported in part by a Discovery Grant from the Natural Sciences and Engineering Research Council of Canada (NSERC) to MJB and by the Intramural Research Program of the NIH and NIAID to MEG. MJB gratefully acknowledges the Canada Research Chair program for salary support.

## Author contributions

MJB, MEG, VP, PS and MLP conceived of the study and planned the experiments. VP, MLP, PS and RR collected the data and all authors were involved in data interpretation. MJB, MEG, PS, VP and RR wrote the manuscript with editorial support from MLP.

## FOOTNOTES

^1^The abbreviations used are: SRS, SAG-Related Sequence; SAG, Surface Antigen Glycoprotein; SPG, Sulfated Proteoglycans; GAG, Glycosaminoglycan; GPI, Glycosylphosphatidylinositol

## FIGURE LEGENDS

**Table S1.** Data collection and refinement statistics.

|  |  |
| --- | --- |
| <u>A. Data collection</u> |  |
| Spacegroup | P2<sub>1</sub>2<sub>1</sub>2<sub>1</sub> |
| a, b, c (Å) | 48.13, 59.71, 101.42 |
| α, β, γ (deg.) | 90, 90, 90 |
| Wavelength | 0.9791 |
| Resolution range (Å) | 43.48 - 1.59 (1.62 - 1.59) |
| Measured reflections | 222901 (10375) |
| Unique reflections | 40027 (1927) |
| Redundancy | 5.6 (5.4) |
| Completeness (%) | 100.0 (99.8) |
| *I*/*σ*(*I*) | 12.6 (2.8) |
| R<sub>merge</sub><sup>a</sup> | 0.064 (0.490) |
| <u>B. Refinement Statistics</u> |  |
| Resolution (Å) | 37.48 - 1.59 |
| R<sub>work</sub><sup>b</sup> / R<sub>free</sub><sup>c</sup> | 0.169/0.191 |
| No. of atoms |  |
| Protein | 2193 |
| Solvent | 325 |
| Glycerol | 6 |
| B-values (Å<sup>2</sup>) |  |
| Protein | 21.8 |
| Solvent | 31.5 |
| Glycerol | 29.8 |
| r.m.s. deviation from ideality |  |
| Bond lengths (Å) | 0.008 |
| Bond angles (deg.) | 1.28 |
| Ramachandran statistics (%) |  |
| Most favoured | 97.4 |
| Allowed | 2.6 |
| Disallowed | 0.0 |

Values in parentheses are for the highest resolution shell
<sup>a</sup> R<sub>merge</sub> = $\sum_{hkl} \sum_i |I_{hkl,i} - [I_{hkl}]| / \sum_{hkl} \sum_i I_{hkl,i}$ , where $[I_{hkl}]$ is the average of symmetry related observations of a unique reflection
<sup>b</sup> R<sub>work</sub> = $\sum |F_{obs} - F_{calc}| / \sum F_{obs}$ , where $F_{obs}$ and $F_{calc}$ are the observed and the calculated structure factors, respectively.
<sup>c</sup> R<sub>free</sub> is R using 5% of reflections randomly chosen and omitted from refinement

**Figure S1:**
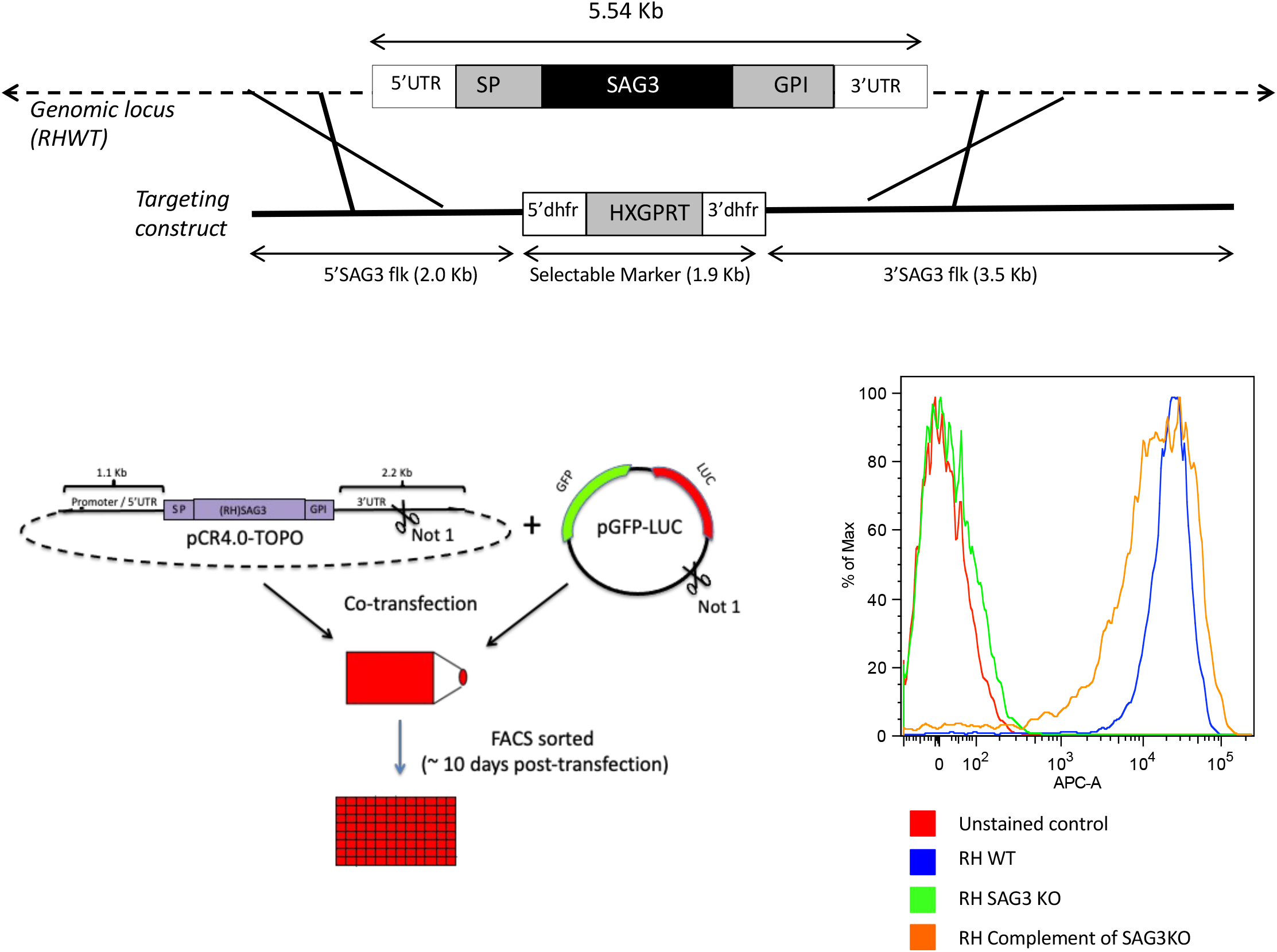
Schematic representation of the strategy for Δ*srs57* complementation and expression of GFP-LUC. 10^7^ RH Δ*srs57* and French RH Δ*srs57* (CL55) parasites were co-transfected with a 3.7 kb PCR amplified genomic locus of Tg*SRS57*, including the promoter and the 5’ and 3’ untranslated regions plus a pGFP-LUC plasmid, containing cDNA sequences of the green fluorescence protein (GFP) and firefly luciferase protein (LUC) for *in vivo* imaging experiments. ∼10 days post-transfection, stable parasites expressing GFP were FACS sorted onto 96 well plates. Positive clones expressing GFP were analyzed for TgSRS57 surface expression by flow cytometry.

**Figure S2:**
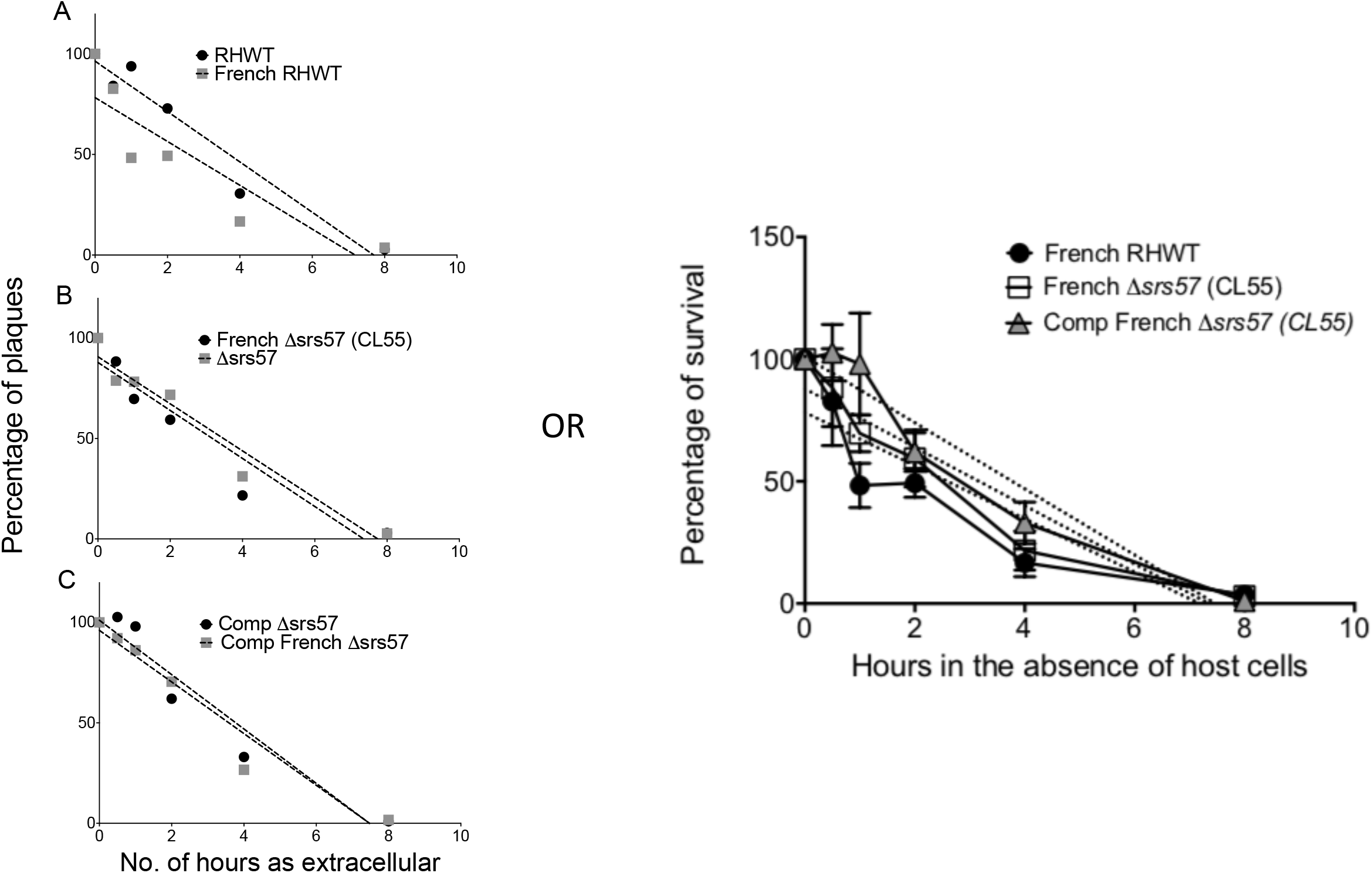
Extracellular survival of French strains. 10^3^ tachyzoites of each strain (French RH WT, French Δ*srs57* and complement) were maintained for 0 min, 30 min, 1, 2, 4 and 8 hrs. in extracellular conditions, before applying to a fresh monolayer of HFF cells. After 5 days, the number of plaques formed at each time point were counted. No significant differences were observed in the extracellular survival and plating efficiency between the strains. Values represent the mean of triplicates ± SD.

**Figure S3:**
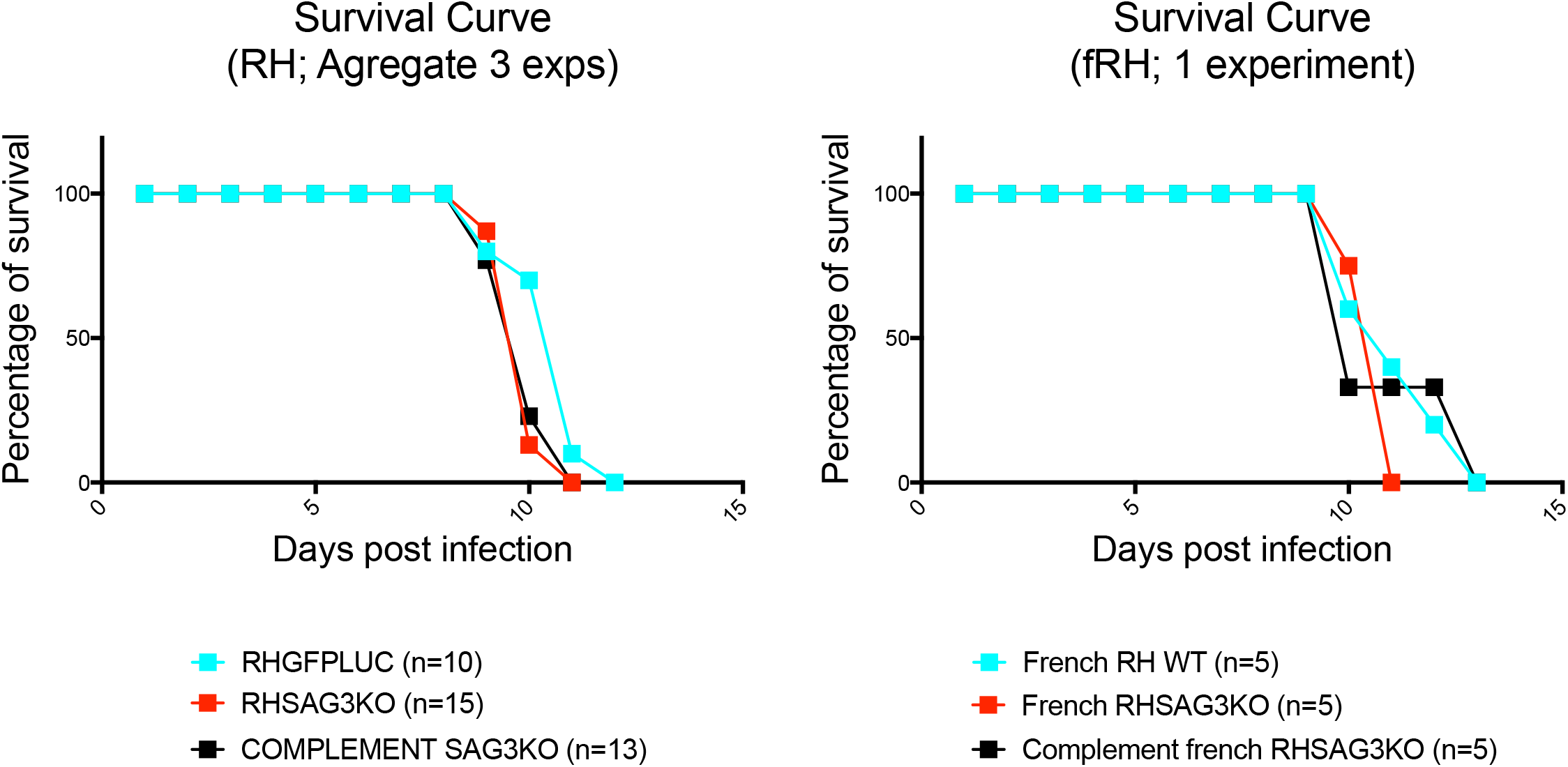
Comparison of mouse survival curves for RH WT, Δ*srs57*, and complemented strains versus French RH WT, Δ*srs57*, and complemented strains. Survival curve of CD-1 mice challenged with 50 tachyzoites/mouse of the indicated strains. No significant differences were observed between the two comparative groups of 3 strains each.

## REFERENCES

1. Tenter AM, Heckeroth AR, Weiss LM. Toxoplasma gondii: from animals to humans. Int J Parasitol. 2000;30(12-13):1217–58. PubMed PMID: 11113252; PubMed Central PMCID: PMCPMC3109627.

2. Grigg ME, Ganatra J, Boothroyd JC, Margolis TP. Unusual abundance of atypical strains associated with human ocular toxoplasmosis. J Infect Dis. 2001;184(5):633–9. PubMed PMID: 11474426.

3. Grigg ME, Dubey JP, Nussenblatt RB. Ocular toxoplasmosis: lessons from Brazil. Am J Ophthalmol. 2015;159(6):999–1001. doi: 10.1016/j.ajo.2015.04.005. PubMed PMID: 25956461; PubMed Central PMCID: PMCPMC4894827.

4. Luft BJ, Remington JS. Toxoplasmic encephalitis in AIDS. Clin Infect Dis. 1992;15(2):211–22.

5. Ancsin JB, Kisilevsky R. A binding site for highly sulfated heparan sulfate is identified in the N terminus of the circumsporozoite protein: significance for malarial sporozoite attachment to hepatocytes. J Biol Chem. 2004;279(21):21824–32. doi: 10.1074/jbc.M401979200. PubMed PMID: 15007056.

6. Carruthers VB, Hakansson S, Giddings OK, Sibley LD. Toxoplasma gondii uses sulfated proteoglycans for substrate and host cell attachment. Infect Immun. 2000;68(7):4005–11. PubMed PMID: 10858215; PubMed Central PMCID: PMCPMC101681.

7. Boothroyd JC, Dubremetz JF. Kiss and spit: the dual roles of Toxoplasma rhoptries. Nat Rev Microbiol. 2008;6(1):79–88. doi: 10.1038/nrmicro1800. PubMed PMID: 18059289.

8. Rostand KS, Esko JD. Microbial adherence to and invasion through proteoglycans. Infect Immun. 1997;65(1):1–8. PubMed PMID: 8975885; PubMed Central PMCID: PMCPMC174549.

9. Bishop JR, Crawford BE, Esko JD. Cell surface heparan sulfate promotes replication of Toxoplasma gondii. Infect Immun. 2005;73(9):5395–401. doi: 10.1128/IAI.73.9.5395-5401.2005. PubMed PMID: 16113255; PubMed Central PMCID: PMCPMC1231081.

10. Bishop JR, Esko JD. The elusive role of heparan sulfate in Toxoplasma gondii infection. Mol Biochem Parasitol. 2005;139(2):267–9. doi: 10.1016/j.molbiopara.2004.11.007. PubMed PMID: 15664661.

11. Ortega-Barria E, Boothroyd JC. A Toxoplasma lectin-like activity specific for sulfated polysaccharides is involved in host cell infection. J Biol Chem. 1999;274(3):1267–76. PubMed PMID: 9880495.

12. Tomley FM, Soldati DS. Mix and match modules: structure and function of microneme proteins in apicomplexan parasites. Trends Parasitol. 2001;17(2):81–8. PubMed PMID: 11228014.

13. Sardinha-Silva A, Mendonça-Natividade FC, Pinzan CF, Lopes CD, Costa DL, Jacot D, et al. The lectin-specific activity of Toxoplasma gondii microneme proteins 1 and 4 binds Toll-like receptor 2 and 4 N-glycans to regulate innate immune priming. PLoS Pathog. 2019;15(6):e1007871. Epub 2019/06/22. doi: 10.1371/journal.ppat.1007871. PubMed PMID: 31226171; PubMed Central PMCID: PMCPMC6608980.

14. Manger ID, Hehl AB, Boothroyd JC. The surface of Toxoplasma tachyzoites is dominated by a family of glycosylphosphatidylinositol-anchored antigens related to SAG1. Infect Immun. 1998;66(5):2237–44. PubMed PMID: 9573113.

15. Jung C, Lee CY-F, Grigg ME. The SRS superfamily of Toxoplasma surface proteins. Intl J Parasitol. 2004;34(1):285–96.

16. Wasmuth JD, Pszenny V, Haile S, Jansen EM, Gast AT, Sher A, et al. Integrated bioinformatic and targeted deletion analyses of the SRS gene superfamily identify SRS29C as a negative regulator of Toxoplasma virulence. MBio. 2012;3(6). doi: 10.1128/mBio.00321-12. PubMed PMID: 23149485; PubMed Central PMCID: PMCPMC3509429.

17. Boulanger MJ, Tonkin ML, Crawford J. Apicomplexan parasite adhesins: novel strategies for targeting host cell carbohydrates. Curr Opin Struct Biol. 2010;20(5):551–9. doi: 10.1016/j.sbi.2010.08.003. PubMed PMID: 20843678.

18. Crawford J, Grujic O, Bruic E, Czjzek M, Grigg ME, Boulanger MJ. Structural characterization of the bradyzoite surface antigen (BSR4) from Toxoplasma gondii, a unique addition to the surface antigen glycoprotein 1-related superfamily. J Biol Chem. 2009;284(14):9192–8. doi: 10.1074/jbc.M808714200. PubMed PMID: 19155215; PubMed Central PMCID: PMCPMC2666571.

19. Reid AJ, Vermont SJ, Cotton JA, Harris D, Hill-Cawthorne GA, Konen-Waisman S, et al. Comparative genomics of the apicomplexan parasites Toxoplasma gondii and Neospora caninum: Coccidia differing in host range and transmission strategy. PLoS Pathog. 2012;8(3):e1002567. doi: 10.1371/journal.ppat.1002567. PubMed PMID: 22457617; PubMed Central PMCID: PMCPMC3310773.

20. Arredondo SA, Cai M, Takayama Y, MacDonald NJ, Anderson DE, Aravind L, et al. Structure of the Plasmodium 6-cysteine s48/45 domain. Proc Natl Acad Sci U S A. 2012;109(17):6692–7. doi: 10.1073/pnas.1204363109. PubMed PMID: 22493233; PubMed Central PMCID: PMCPMC3340019.

21. Gerloff DL, Creasey A, Maslau S, Carter R. Structural models for the protein family characterized by gamete surface protein Pfs230 of Plasmodium falciparum. Proc Natl Acad Sci U S A. 2005;102(38):13598–603. doi: 10.1073/pnas.0502378102. PubMed PMID: 16155126; PubMed Central PMCID: PMCPMC1224620.

22. Kennedy AT, Schmidt CQ, Thompson JK, Weiss GE, Taechalertpaisarn T, Gilson PR, et al. Recruitment of Factor H as a Novel Complement Evasion Strategy for Blood-Stage Plasmodium falciparum Infection. J Immunol. 2016;196(3):1239–48. doi: 10.4049/jimmunol.1501581. PubMed PMID: 26700768.

23. Parker ML, Peng F, Boulanger MJ. The Structure of Plasmodium falciparum Blood-Stage 6-Cys Protein Pf41 Reveals an Unexpected Intra-Domain Insertion Required for Pf12 Coordination. PLoS One. 2015;10(9):e0139407. doi: 10.1371/journal.pone.0139407. PubMed PMID: 26414347; PubMed Central PMCID: PMCPMC4587554.

24. Tonkin ML, Arredondo SA, Loveless BC, Serpa JJ, Makepeace KA, Sundar N, et al. Structural and biochemical characterization of Plasmodium falciparum 12 (Pf12) reveals a unique interdomain organization and the potential for an antiparallel arrangement with Pf41. J Biol Chem. 2013;288(18):12805–17. doi: 10.1074/jbc.M113.455667. PubMed PMID: 23511632; PubMed Central PMCID: PMCPMC3642325.

25. Azzouz N, Kamena F, Laurino P, Kikkeri R, Mercier C, Cesbron-Delauw MF, et al. Toxoplasma gondii secretory proteins bind to sulfated heparin structures. Glycobiology. 2013;23(1):106–20. doi: 10.1093/glycob/cws134. PubMed PMID: 22997241.

26. Jacquet A, Coulon L, De Neve J, Daminet V, Haumont M, Garcia L, et al. The surface antigen SAG3 mediates the attachment of Toxoplasma gondii to cell-surface proteoglycans. Mol Biochem Parasitol. 2001;116(1):35–44. PubMed PMID: 11463464.

27. Mineo JR, McLeod R, Mack D, Smith J, Khan IA, Ely KH, et al. Antibodies to Toxoplasma gondii major surface protein (SAG-1, P30) inhibit infection of host cells and are produced in murine intestine after peroral infection. J Immunol. 1993;150(9):3951–64.

28. Dzierszinski F, Mortuaire M, Cesbron-Delauw MF, Tomavo S. Targeted disruption of the glycosylphosphatidylinositol-anchored surface antigen SAG3 gene in Toxoplasma gondii decreases host cell adhesion and drastically reduces virulence in mice. Mol Microbiol. 2000;37(3):574–82. PubMed PMID: 10931351.

29. He XL, Grigg ME, Boothroyd JC, Garcia KC. Structure of the immunodominant surface antigen from the Toxoplasma gondii SRS superfamily. Nat Struct Biol. 2002;9(8):606–11. PubMed PMID: 12091874.

30. Fuhrman SA, Joiner KA. Toxoplasma gondii: mechanism of resistance to complement-mediated killing. J Immunol. 1989;142(3):940–7. Epub 1989/02/01. PubMed PMID: 2643665.

31. Sikorski PM, Commodaro AG, Grigg ME. Toxoplasma gondii Recruits Factor H and C4b-Binding Protein to Mediate Resistance to Serum Killing and Promote Parasite Persistence in vivo. Front Immunol. 2019;10:3105. Epub 2020/02/06. doi: 10.3389/fimmu.2019.03105. PubMed PMID: 32010145; PubMed Central PMCID: PMCPMC6979546.

32. Sikorski PM, Commodaro AG, Grigg ME. A Protective and Pathogenic Role for Complement During Acute Toxoplasma gondii Infection. Front Cell Infect Microbiol. 2021;11:634610. Epub 2021/03/12. doi: 10.3389/fcimb.2021.634610. PubMed PMID: 33692968; PubMed Central PMCID: PMCPMC7937796.

33. Blaum BS. The lectin self of complement factor H. Curr Opin Struct Biol. 2017;44:111–8. Epub 2017/02/13. doi: 10.1016/j.sbi.2017.01.005. PubMed PMID: 28189794.

34. Kajander T, Lehtinen MJ, Hyvarinen S, Bhattacharjee A, Leung E, Isenman DE, et al. Dual interaction of factor H with C3d and glycosaminoglycans in host-nonhost discrimination by complement. Proc Natl Acad Sci U S A. 2011;108(7):2897–902. Epub 2011/02/03. doi: 10.1073/pnas.1017087108. PubMed PMID: 21285368; PubMed Central PMCID: PMCPMC3041134.

35. Crawford J, Lamb E, Wasmuth J, Grujic O, Grigg ME, Boulanger MJ. Structural and functional characterization of SporoSAG: a SAG2-related surface antigen from Toxoplasma gondii. J Biol Chem. 2010;285(16):12063–70. doi: 10.1074/jbc.M109.054866. PubMed PMID: 20164173; PubMed Central PMCID: PMCPMC2852944.

36. Fadel S, Eley A. Chlorate: a reversible inhibitor of proteoglycan sulphation in Chlamydia trachomatis-infected cells. J Med Microbiol. 2004;53(Pt 2):93–5. doi: 10.1099/jmm.0.05497-0. PubMed PMID: 14729927.

37. Humphries DE, Silbert JE. Chlorate: a reversible inhibitor of proteoglycan sulfation. Biochem Biophys Res Commun. 1988;154(1):365–71. PubMed PMID: 2969240.

38. Gavrilescu LC, Denkers EY. IFN-gamma overproduction and high level apoptosis are associated with high but not low virulence Toxoplasma gondii infection. J Immunol. 2001;167(2):902–9. PubMed PMID: 11441097.

39. Mordue DG, Monroy F, La Regina M, Dinarello CA, Sibley LD. Acute toxoplasmosis leads to lethal overproduction of Th1 cytokines. J Immunol. 2001;167(8):4574–84. PubMed PMID: 11591786.

40. Couper KN, Roberts CW, Brombacher F, Alexander J, Johnson LL. Toxoplasma gondii-specific immunoglobulin M limits parasite dissemination by preventing host cell invasion. Infect Immun. 2005;73(12):8060–8. Epub 2005/11/22. doi: 10.1128/IAI.73.12.8060-8068.2005. PubMed PMID: 16299300; PubMed Central PMCID: PMCPMC1307022.

41. Aquino RS, Park PW. Glycosaminoglycans and infection. Front Biosci (Landmark Ed). 2016;21:1260–77. PubMed PMID: 27100505; PubMed Central PMCID: PMCPMC4975577.

42. Esko JD, Lindahl U. Molecular diversity of heparan sulfate. J Clin Invest. 2001;108(2):169–73. doi: 10.1172/JCI13530. PubMed PMID: 11457867; PubMed Central PMCID: PMCPMC203033.

43. Rogerson SJ, Chaiyaroj SC, Ng K, Reeder JC, Brown GV. Chondroitin sulfate A is a cell surface receptor for Plasmodium falciparum-infected erythrocytes. J Exp Med. 1995;182(1):15–20. PubMed PMID: 7790815; PubMed Central PMCID: PMCPMC2192085.

44. Rieger H, Yoshikawa HY, Quadt K, Nielsen MA, Sanchez CP, Salanti A, et al. Cytoadhesion of Plasmodium falciparum-infected erythrocytes to chondroitin-4-sulfate is cooperative and shear enhanced. Blood. 2015;125(2):383–91. doi: 10.1182/blood-2014-03-561019. PubMed PMID: 25352129; PubMed Central PMCID: PMCPMC4287643.

45. Coppi A, Tewari R, Bishop JR, Bennett BL, Lawrence R, Esko JD, et al. Heparan sulfate proteoglycans provide a signal to Plasmodium sporozoites to stop migrating and productively invade host cells. Cell Host Microbe. 2007;2(5):316–27. doi: 10.1016/j.chom.2007.10.002. PubMed PMID: 18005753; PubMed Central PMCID: PMCPMC2117360.

46. Robert R, de la Jarrige PL, Mahaza C, Cottin J, Marot-Leblond A, Senet JM. Specific binding of neoglycoproteins to Toxoplasma gondii tachyzoites. Infect Immun. 1991;59(12):4670–3. PubMed PMID: 1937826; PubMed Central PMCID: PMCPMC259094.

47. van Dijk MR, van Schaijk BC, Khan SM, van Dooren MW, Ramesar J, Kaczanowski S, et al. Three members of the 6-cys protein family of Plasmodium play a role in gamete fertility. PLoS Pathog. 2010;6(4):e1000853. doi: 10.1371/journal.ppat.1000853. PubMed PMID: 20386715; PubMed Central PMCID: PMCPMC2851734.

48. Aslam M, Perkins SJ. Folded-back solution structure of monomeric factor H of human complement by synchrotron X-ray and neutron scattering, analytical ultracentrifugation and constrained molecular modelling. J Mol Biol. 2001;309(5):1117–38. Epub 2001/06/12. doi: 10.1006/jmbi.2001.4720. PubMed PMID: 11399083.

49. Powell AK, Yates EA, Fernig DG, Turnbull JE. Interactions of heparin/heparan sulfate with proteins: appraisal of structural factors and experimental approaches. Glycobiology. 2004;14(4):17R–30R. doi: 10.1093/glycob/cwh051. PubMed PMID: 14718374.

50. Lacy HM, Sanderson RD. 6xHis promotes binding of a recombinant protein to heparan sulfate. Biotechniques. 2002;32(2):254, 6, 8. PubMed PMID: 11848397.

51. McLeod R, Skamene E, Brown CR, Eisenhauer PB, Mack DG. Genetic regulation of early survival and cyst number after peroral Toxoplasma gondii infection of A x B/B x A recombinant inbred and B10 congenic mice. J Immunol. 1989;143(9):3031–4. PubMed PMID: 2809214.

52. Joiner KA, Fuhrman SA, Miettinen HM, Kasper LH, Mellman I. Toxoplasma gondii: fusion competence of parasitophorous vacuoles in Fc receptor-transfected fibroblasts. Science. 1990;249(4969):641–6. Epub 1990/08/10. doi: 10.1126/science.2200126. PubMed PMID: 2200126.

53. Zhao Y, Marple AH, Ferguson DJ, Bzik DJ, Yap GS. Avirulent strains of Toxoplasma gondii infect macrophages by active invasion from the phagosome. Proc Natl Acad Sci U S A. 2014;111(17):6437–42. Epub 2014/04/16. doi: 10.1073/pnas.1316841111. PubMed PMID: 24733931; PubMed Central PMCID: PMCPMC4035997.

54. Han C, Jin J, Xu S, Liu H, Li N, Cao X. Integrin CD11b negatively regulates TLR-triggered inflammatory responses by activating Syk and promoting degradation of MyD88 and TRIF via Cbl-b. Nat Immunol. 2010;11(8):734–42. Epub 2010/07/20. doi: 10.1038/ni.1908. PubMed PMID: 20639876.

55. Zhang X, Kimura Y, Fang C, Zhou L, Sfyroera G, Lambris JD, et al. Regulation of Toll-like receptor-mediated inflammatory response by complement in vivo. Blood. 2007;110(1):228–36. Epub 2007/03/17. doi: 10.1182/blood-2006-12-063636. PubMed PMID: 17363730; PubMed Central PMCID: PMCPMC1896115.

56. Huang JH, Lin CY, Wu SY, Chen WY, Chu CL, Brown GD, et al. CR3 and Dectin-1 Collaborate in Macrophage Cytokine Response through Association on Lipid Rafts and Activation of Syk-JNK-AP-1 Pathway. PLoS Pathog. 2015;11(7):e1004985. Epub 2015/07/02. doi: 10.1371/journal.ppat.1004985. PubMed PMID: 26132276; PubMed Central PMCID: PMCPMC4488469.

57. Hajishengallis G, Lambris JD. More than complementing Tolls: complement-Toll-like receptor synergy and crosstalk in innate immunity and inflammation. Immunol Rev. 2016;274(1):233–44. Epub 2016/10/27. doi: 10.1111/imr.12467. PubMed PMID: 27782328; PubMed Central PMCID: PMCPMC5119927.

58. Liszewski MK, Kolev M, Le Friec G, Leung M, Bertram PG, Fara AF, et al. Intracellular complement activation sustains T cell homeostasis and mediates effector differentiation. Immunity. 2013;39(6):1143–57. Epub 2013/12/10. doi: 10.1016/j.immuni.2013.10.018. PubMed PMID: 24315997; PubMed Central PMCID: PMCPMC3865363.

59. Strainic MG, Liu J, Huang D, An F, Lalli PN, Muqim N, et al. Locally produced complement fragments C5a and C3a provide both costimulatory and survival signals to naive CD4+ T cells. Immunity. 2008;28(3):425–35. Epub 2008/03/11. doi: 10.1016/j.immuni.2008.02.001. PubMed PMID: 18328742; PubMed Central PMCID: PMCPMC2646383.

60. Carroll MC. Complement and humoral immunity. Vaccine. 2008;26 Suppl 8:I28–33. Epub 2009/04/24. doi: 10.1016/j.vaccine.2008.11.022. PubMed PMID: 19388161; PubMed Central PMCID: PMCPMC4018718.

61. Dempsey PW, Allison ME, Akkaraju S, Goodnow CC, Fearon DT. C3d of complement as a molecular adjuvant: bridging innate and acquired immunity. Science. 1996;271(5247):348–50. Epub 1996/01/19. doi: 10.1126/science.271.5247.348. PubMed PMID: 8553069.

62. Leslie AGW. Recent changes to the MOSFLM package for processing film and image plate data. Joint CCP4 + ESF-EAMCB Newsletter on Protein Crystallography. 1992;26.

63. Evans P. Scaling and assessment of data quality. Acta Crystallogr D Biol Crystallogr. 2006;62(Pt 1):72–82. doi: 10.1107/S0907444905036693. PubMed PMID: 16369096.

64. Collaborative CP. The CCP4 suite: programs for protein crystallography. Acta Crystallogr D Biol Crystallogr. 1994;50:760–3.

65. McCoy AJ, Grosse-Kunstleve RW, Adams PD, Winn MD, Storoni LC, Read RJ. Phaser crystallographic software. J Appl Crystallogr. 2007;40(Pt 4):658–74. doi: 10.1107/S0021889807021206. PubMed PMID: 19461840; PubMed Central PMCID: PMCPMC2483472.

66. Stein N. CHAINSAW: a program for mutating pdb files used as templates in molecular replacement. J Appl Crystallogr. 2008;41:641–3.

67. Emsley P, Cowtan K. Coot: model-building tools for molecular graphics. Acta Crystallogr D Biol Crystallogr. 2004;60(Pt 12 Pt 1):2126–32. doi: 10.1107/S0907444904019158. PubMed PMID: 15572765.

68. Murshudov GN, Skubak P, Lebedev AA, Pannu NS, Steiner RA, Nicholls RA, et al. REFMAC5 for the refinement of macromolecular crystal structures. Acta Crystallogr D Biol Crystallogr. 2011;67(Pt 4):355–67. doi: 10.1107/S0907444911001314. PubMed PMID: 21460454; PubMed Central PMCID: PMCPMC3069751.

69. Baker NA, Sept D, Joseph S, Holst MJ, McCammon JA. Electrostatics of nanosystems: application to microtubules and the ribosome. Proc Natl Acad Sci U S A. 2001;98(18):10037–41. doi: 10.1073/pnas.181342398. PubMed PMID: 11517324; PubMed Central PMCID: PMCPMC56910.

